# Homoplastic evolution of proximal podomere patterning across Arthropoda

**DOI:** 10.64898/2026.10.30.763126

**Authors:** Benjamin C. Klementz, Max Hämmerle, Ethan M. Laumer, Sophie M. Neu, Madeline M. Koebrick, Nithya Atla, Pranil Panda, Neeharika Verma, Georg Brenneis, Prashant P. Sharma

## Abstract

The jointed arthropod leg is often credited as a key innovation spurring the evolutionary success of the phylum. Across lineages, each territory of the appendage has served as the substrate for myriad adaptations and structural innovations. While much of the available data has supported a conserved suite of transcription factors responsible for the initiation and patterning of the arthropod appendage, recent works have begun to highlight instances of lineage-specific shifts in expression dynamics, particularly amongst the chelicerates, correlated both with the emergence of novel structures and developmental system drift in the specification of plesiomorphic structures. These works confound efforts to align or homologize the leg segments across arthropod lineages, an effort already challenged by ground plan differences in segment number between the subphyla. An even more unusual architecture of leg segments occurs in sea spiders (Pycnogonida), which alone among Arthropoda are considered by systematists to bear three coxae (proximal segments), in contrast to the single coxa of the remaining arthropods. Here, we examined proximal podomere patterning in the developing appendages of a sea spider and an arachnid. We find that a particular spatial arrangement of *Lim1* and *SoxNeuro* characterizes the developing coxa and trochanter of the harvestman *Phalangium opilio* (Opiliones) throughout appendage formation, with RNAi depletions of *Lim1* yielding deletions of coxa and trochanter. Comparison of expression patterns across taxa shows that *Lim1* and *SoxNeuro* have a similar spatial arrangement in the harvestman and sea spider, but these suggest conflicting interpretations of leg segment homology. We show that this is attributable to homoplastic patterns of evolution in *SoxNeuro* across Arthropoda. Expanding our surveys of prominent PD axis-patterning transcription factors supports additional instances of homoplasy in the domains of *Sp6-9* and *Irx2*. The pervasiveness of homoplasy in gene expression patterns demands measured skepticism of homology statements and careful comparisons of multiple patterning genes, toward achieving falsifiability in the homology of structures.

## Introduction

The modularity of the arthropod body plan, facilitated by the division of the body into segmental units, has long been one of many explanations for the phylum’s evolutionary success and high level of extant diversity, alongside key innovations such as the wings of insects or silks of spiders (Bond and Opell, 1998; Dudley and Pass, 2018; Stansbury and Moczek, 2013). This modularity exists along two major axes of the arthropod body. Most prominent is the division of the primary—or anteroposterior (AP)—body axis into discrete segments. This division, in turn, was secondarily modified into lineage-specific patterns of tagmosis, fusions of adjacent segments into functionally specialized body regions, with particular patterns often diagnostic of each of the four subphyla. Such functional specialization is achieved principally by variation in the architecture of the paired segmental appendages, typified by the morphological disparity between the feeding mandibles of the insect head and the thoracic walking legs (Adamowicz et al., 2008). Gross changes in appendage identity are governed by the Hox genes, which confer segmental identities along the AP axis (Akam, 1995; Gainett et al., 2023; Lewis, 1978; Martin et al., 2016). Division of the proximodistal (PD) appendage axis into segments is the secondary axis upon which modularity facilitates body plan disparity and functional specialization. Variations in the number, size, and shape of individual appendage segments, or podomeres, can achieve myriad functional outcomes. As examples, the subdivision of the tarsus into numerous tarsomeres in lineages like Eupnoi harvestmen confer prehensility, while in orthopterans, hypertrophied femora and tibiae of the T3 leg pair house the musculature required for their remarkable jumping ability.

The developmental genetic mechanisms underlying the evolution of arthropod appendages are now informed by functional datasets spanning both mandibulate and chelicerate model species. Broadly, comparative assessments of PD axis patterning have supported conserved functions of many transcription factors, in particular the cofactors *homothorax* and *extradenticle, dachshund*, and *Distal-less* in the regionalization of proximal, medial, and distal appendage territories, respectively (Abzhanov and Kaufman, 2000; Dong et al., 2001; Panganiban et al., 1997, 1995, 1994). Recent works in chelicerates, however, have noted several instances of lineage-specific divergences in the function of PD axis transcription factors (Klementz et al., 2026, 2024). The homeobox gene *clawless* (*C15*) is required for the formation of the pretarsus and its claws in insects, suppressing the tarsal patterning gene *Bar* in the distal appendage terminus. Yet in several chelicerates, *clawless* is not expressed in the pretarsus during stages when claw formation occurs. Instead, *clawless* is expressed at the metatarsus-tarsus border, with functional experiments confirming a role in maintaining this boundary (Klementz et al., 2026). Other experiments have documented lineage-specific gains or losses of podomeres as a result of divergence in transcription factor expression domains. As an example, while *dachshund* expression in the proximal podomere of the harvestman chelicera maintains the ancestral three-segmented architecture, *dachshund* is absent from the derived, two-segmented chelicerae of spiders (Prpic and Damen, 2004; Sharma et al., 2013). Likewise, a distal ring of *extradenticle* in the medial territory of the chelicerate pedipalps and walking legs underlies the origin of the patella (Klementz et al., 2024). Despite lacking functional validation, recent expression surveys have also observed several other cases of divergent transcription factor expression domains along the PD axis of chelicerates in comparison to insect models, including genes such as *SoxNeuro, apterous, BarH1, bric-a-brac, trachealess, rotund*, and *Lim1* (Klementz et al., 2025; Zhang, 2016). Even within lineages, the expression and function of these transcription factors in appendage formation can be highly variable, exemplified by dissimilar podomere fusions following *nubbin* RNAi in several insect lineages, and shifts in *homothorax* expression across spider species (Turchyn et al., 2011; Turetzek et al., 2017).

Such changes confound inferences of podomere homologies across arthropods, a challenge already exacerbated by lineage-specific patterns of appendage segmentation (Fig. 1). While insects invariably possess a five-segmented walking leg (coxa, trochanter, femur, tibia, tarsus), chelicerate legs are most often composed of six podomeres (coxa, trochanter, femur, patella, tibia, and tarsus). The metatarsus of many arachnids is variably interpreted as either a true podomere or a subdivision of the ancestral tarsus. Here, we treat the metatarsus as a tarsal article to facilitate direct comparison of podomere homologies with a prior work (Bruce, 2021). In both insects and arachnids, an additional element, the pretarsus, occurs at the distal appendage terminus bearing the claws. Morphological changes to individual podomeres across chelicerates have led to variable inferences of subdivided trochanters or femora in lineages such as solifuges or hooded tickspiders (reviewed in Klementz et al., 2025). Yet, the largest disparity in appendage architecture occurs in the sea spiders (Pycnogonida). The leg of the pycnogonid crown-group is often considered to consist of eight podomeres. The three proximal most podomeres, due to their smaller size and position, are believed to represent three coxal homologs, followed by femur, patella (alternatively, tibia 1), tibia (alternatively, tibia 2), tarsus, and propodus which bears the main claw (Fig. 1A-B). As with the metatarsus, we hereafter consider the sea spider tarsus and propodus to represent similar subdivisions of a single segment to align with prior inferences (Bruce, 2021). The prominent main claw is also considered a podomere, similar to the often claw-shaped crustacean dactylus or the tarsus of crown-group xiphosurans (e.g., Brenneis and Sholtz, 2021; Bruce, 2021). The structure of the palps and ovigers is even more unusual and variable, displaying in some families as many as 9 or 10 distinct articles, respectively (Arango and Wheeler, 2007; Bamber, 2010; Cano-Sánchez and López-González, 2016). Fossilized specimens may also bear additional podomeres in the legs, palps, and ovigers (Sabroux et al., 2024).

**Fig. 1.**
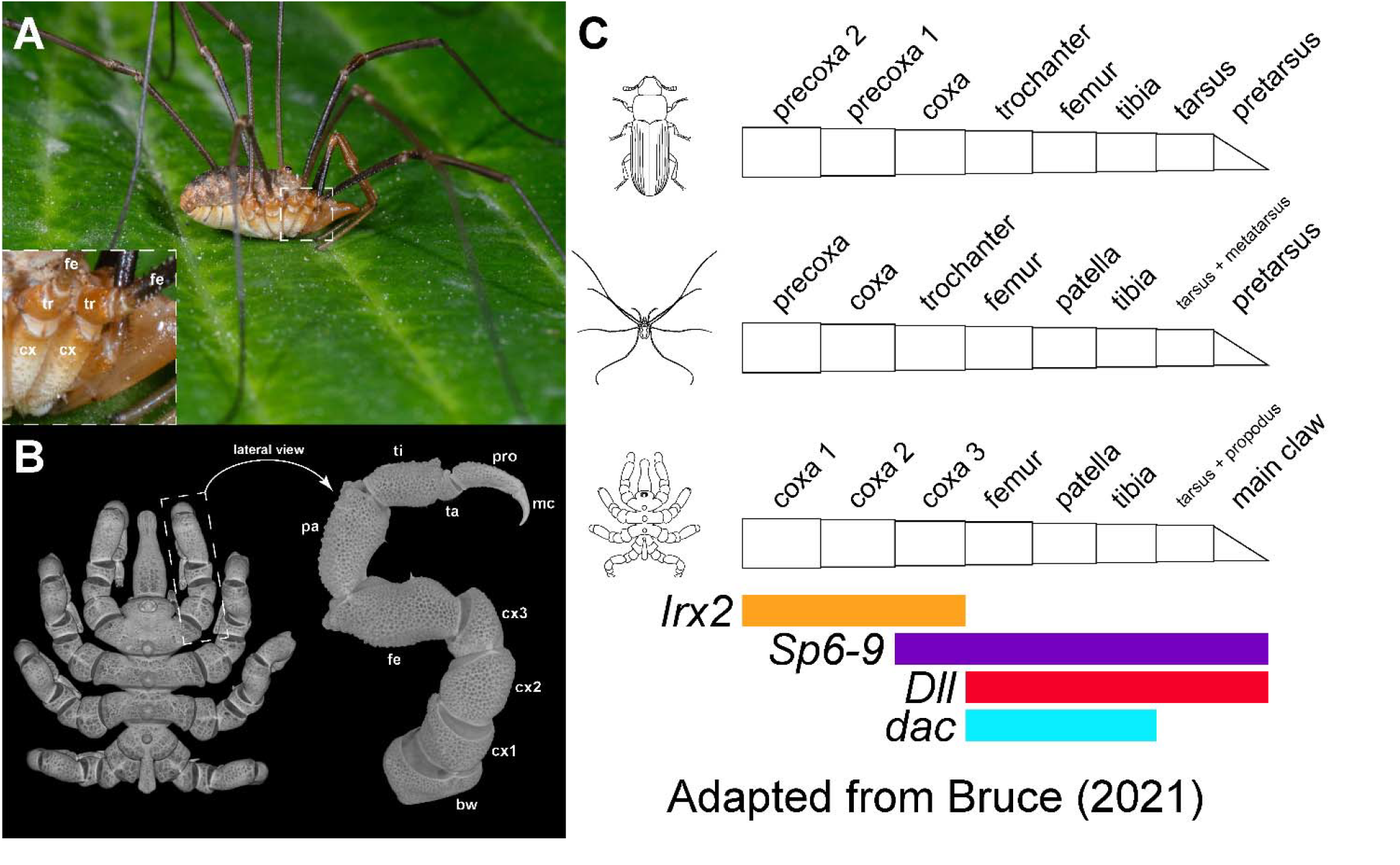
Variance in chelicerate proximal leg segmentation and the alignment of the arthropod PD axis. (A) Live habitus of the harvestman *Phalangium opilio* in lateral view. Inset: magnification of leg I and II proximal appendage podomeres. Note the presence of a singular coxa and trochanter. (B) Fixed specimen of the sea spider *Pycnogonum litorale* in dorsal view (left) with magnification of walking leg I in lateral view (right). Note the presence of three coxae and absence of a trochanter. (C) Hypothesized alignment of podomeres between insect and chelicerate exemplars adapted from Bruce (2021). Colored bars beneath the appendages represent simplified depictions of otherwise heterogeneous expression domains of prominent PD axis transcription factors leveraged to support homologies of individual territories across lineages in Bruce (2021). Abbreviations: cx – coxa; tr – trochanter; fe – femur.

While appendage architecture can vary across lineages, similarities in expression dynamics of PD axis-patterning transcription factors between chelicerate lineages, particularly when localized to individual podomeres and functional experiments have validated a role in the maintenance of that territory, have informed novel hypotheses of podomere homologies. Conserved, distal expression of *extradenticle* across several distantly-related chelicerate taxa supports retention of patellar homologs across the subphylum (Klementz et al., 2024). Similarly, shared patterns of *clawless* expression at the metatarsus-tarsus boundary support inferences of homology between the arachnid tarsus and the propodus of sea spiders, despite major differences in the degree of tarsomerization (Klementz et al., 2026). Expanding on these efforts, we recently proposed to homologize the singular coxae of arachnids, with the third coxae of sea spiders (Klementz et al., 2025). This hypothesis was predicated on the observation of a ring of *SoxNeuro* in the distal compartment of the coxa in a harvestman, and comparable expression in the third coxae of sea spiders, suggesting that sea spiders may indeed possess three coxal podomeres. This result was surprising given that *SoxNeuro* is not expressed in the coxae of insects, instead localizing to the embryonic central nervous system, corpus allatum, and developing epidermal denticles (Crémazy et al., 2000; Luo et al., 2026; Rizzo and Bejsovec, 2017). A recent survey of Sox family genes in a spider, however, did not explicitly note appendicular expression of *SoxNeuro* during stages when the gene is active in the embryonic neuroectoderm (Bonatto Paese et al., 2018). As the authors surveyed only early limb bud stages, presence in the appendages is ambiguous.

Our prior expression survey, however, noted another potential candidate for the alignment of proximal leg segments between sea spiders and arachnids, the homeobox gene *Lim1*. In the fruit fly, *Lim1* is expressed in ring domains in the coxa, femur, and tibia, and strongly at the distal terminus of the appendage (Tsuji et al., 2000). In the distal territory, *Lim1* acts cooperatively with *aristaless* to repress *Bar* during the separation of tarsus and pretarsus. Proximally, depletions of *Lim1* expression yield truncations of the femur and loss of the coxae, with additional malformation of the joint between femur and tibia (Pueyo et al., 2000; Tsuji et al., 2000). RNA interference (RNAi) against *Lim1* in the red flour beetle *Tribolium castaneum* recapitulated the same phenotypic outcomes of deleted proximal podomeres and malformed medial joints, suggesting a conserved function of *Lim1* across hexapods (Angelini et al., 2012). The initial survey of *Lim1* expression in the harvestman, despite surveying only a single embryonic stage, suggested similar dynamics, with a strong ring-domain of expression localizing to the distal coxa and trochanter, but lacking the medial expression characteristic of insect models.

Here, to redress the paucity of expression data, we characterize the dynamics of *Lim1* and *SoxNeuro* in the model harvestman, *Phalangium opilio*, throughout stages of appendage formation and elongation. To assess the function of harvestman *Lim1* in proximal leg segmentation, we performed RNAi against this transcription factor in *P. opilio*. We then compared the expression territories of *Lim1* and *SoxNeuro* in the harvestman with the sea spider *Pycnogonum litorale* to infer podomere homologies between both chelicerate lineages. Additionally, we studied the expression dynamics of two other prominent PD axis-patterning transcription factors, *Sp6-9* and *Irx2* (i.e., *araucan*), recently hypothesized to serve as additional markers of proximal podomere identity in the alignment of crustacean and insect appendages, to test for the proposed presence of a cryptic precoxa that was incorporated into the body wall in non-sea spider arthropods (Bruce and Patel, 2020; Bruce, 2021) (Fig. 1C). We show that spatial and functional dynamics of proximal leg patterning genes exhibit a high degree of lability across Arthropoda. Neither chelicerate lineage exhibited evidence of a precoxa homolog. Our results substantiate additional examples of lineage-specific modifications to the appendage segmentation program and homoplastic patterns of evolution in the PD axis-patterning gene regulatory network.

## Materials and Methods

### Orthology Inference

Homologs of *Phalangium opilio Lim1, SoxNeuro, Sp6-9, dachshund, odd-skipped*, and *Irx2*, and *Pycnogonum litorale SoxNeuro* and *dachshund* were identified in previously published works (Klementz et al., 2026, 2025, 2024; Setton et al., 2024; Setton and Sharma, 2018).

All other genes were queried from the genomes or developmental transcriptomes of target species by a series of tBLASTn searches, with identities secondarily validated by SMART-BLAST; NCBI accession details of queried assemblies can be found in Supplementary File S1, Table S1. The single-copy *SoxNeuro* homologs of the centipede *Lithobius atkinsoni* and amphipod *Parhyale hawaiensis* were queried using the peptide sequence of *D. melanogaster* (NP_001260269.1), and the *P. hawaiensis Lim1* homolog was queried using the peptide sequence of *D. melanogaster* (NP_572505.1). The *Pycnogonum litorale Lim1* homolog (PB.6076) was recovered from the *P. litorale* genome (Papadopoulos et al. 2025) using the peptide sequence of *T. castaneum* (EFA05667.1) as a query. The *P. litorale Irx2* ortholog (g10269) was previously identified by Papadopoulos et al. (2025), being therein designated as *Irx1* based on the ancestral syntenic order of *Irx* genes in arthropods (Aase-Remedios et al. 2023).

### Gene Expression Assays

Embryos of *P. opilio* and *P. hawaiensis*, as well as post-embryonic instars of *P. litorale* were cultivated, fixed, and assayed for gene expression via fluorescent hybridization chain reaction (HCR) following previously established protocols (Bruce et al., 2021; Gainett et al., 2021; Klementz et al., 2024). Embryo cultivation and fixation for *L. atkinsoni* followed the protocol of (Setton et al., 2017). Identical protocols to that of *P. opilio* produced comparable expression intensities following HCR in the centipede.

For genes not previously identified, HCR probe pairs were designed with the HCR Probe Maker tool or generated via submission of CDS to Molecular Instruments (Kuehn et al., 2022). Output probe pair sequences from the HCR Probe Maker tool and CDS sequences for proprietary probes designed by Molecular Instruments are both provided in Supplementary File S1 (Supplementary materials and methods; tables S2-S8). Design details for all other genes are available in prior publications (Klementz et al., 2026, 2025a, 2024; Setton et al., 2024).

### RNA Interference

Primer pairs designed for the cloning of *Po-Lim1* are provided in Supplementary File S1 (table S9). *Po-Lim1* fragments were amplified via standard PCR protocols and cloned using a TOPO TA Cloning Kit with One Shot Top10 chemically competent *Escherichia coli* (ThermoFisher) following the manufacturer’s protocol. Sanger sequencing with M13 primers was used to validate product identity. Double-stranded RNA (dsRNA) was synthesized using the MEGAscript T7 kit (Ambion/Life Technologies). dsRNA concentrations following synthesis were assessed using a NanoDrop ONE spectrophotometer before adjustment to final concentrations of 4 µg/µL. Vital dyes were added to final dilutions in a volumetric ratio of 1:20 for visualization during injection. Microinjection was performed on synchronously developing clutches of embryos as previous described (Sharma et al., 2013). Unlike previous analyses of PD appendage axis patterning genes, injections targeting limb bud stages did not produce any phenotypic outcomes. Instead, injections targeted stages following initial appearance of fluid-filled space between the embryonic tissue and the perivitelline membrane (stages 5-6; nomenclature following Gainett et al., 2022). Embryos designated negative control were injected with distilled water mixed with vital dye under otherwise identical conditions. Following injection, both negative control and RNA interference (RNAi) embryos were reared in a 26°C incubator, with subsets fixed for HCR at representative stages of appendage formation and elongation. Those not fixed for HCR assays were left to develop until the first appearance of hatchlings, before manual removal from the vitelline membrane for subsequent brightfield imaging. Hatchlings were repeatedly washed in heptane to remove residual halocarbon oil followed by progressive washes into a final mixture of 75% glycerol and PBS-T.

### Imaging

Brightfield images of *P. opilio* appendages were captured on an Olympus BX60 epifluorescence microscope driven by cellSens software. Appendages were imaged at varying focal planes and compiled into focused stacks with Helicon Focus v. 6.6.1. Details on micro-computed X-ray tomography of adult *P. litorale* are described in a previous study (Frankowski et al., 2022). Confocal laser scanning microscopy of HCR embryos was performed on both Zeiss LSM 780 and Zeiss LSM 980 microscopes driven by Zen software for *P. opilio, P. hawaiensis*, and *L. atkinsoni*. HCR imaging of *P. litorale* was performed with both Leica SP5 and Stellaris 5 microscopy platforms driven by LAS(X)-AF software. Cuticular autofluorescence was documented separately in the channel of the 488 nm laser line and subsequently used to mask the cuticle in the gene expression channels using Amira 3D v. 2021.1 as previously described (Klementz et al., 2024).

## Results

### Strong Po-Lim1 expression marks the developing coxa and trochanter throughout appendage development

Following their initial emergence from the body wall at stage 8 (stages following Gainett et al., 2022), *Po-Lim1* is strongly expressed in a ring domain in the proximal-most territory of each appendage (Fig. 2A). Outside of the paired segmental appendages, *Po-Lim1* is detected in a pair of bilaterally symmetrical patches subtending the head lobes. No expression is detected in the labrum.

**Fig. 2.**
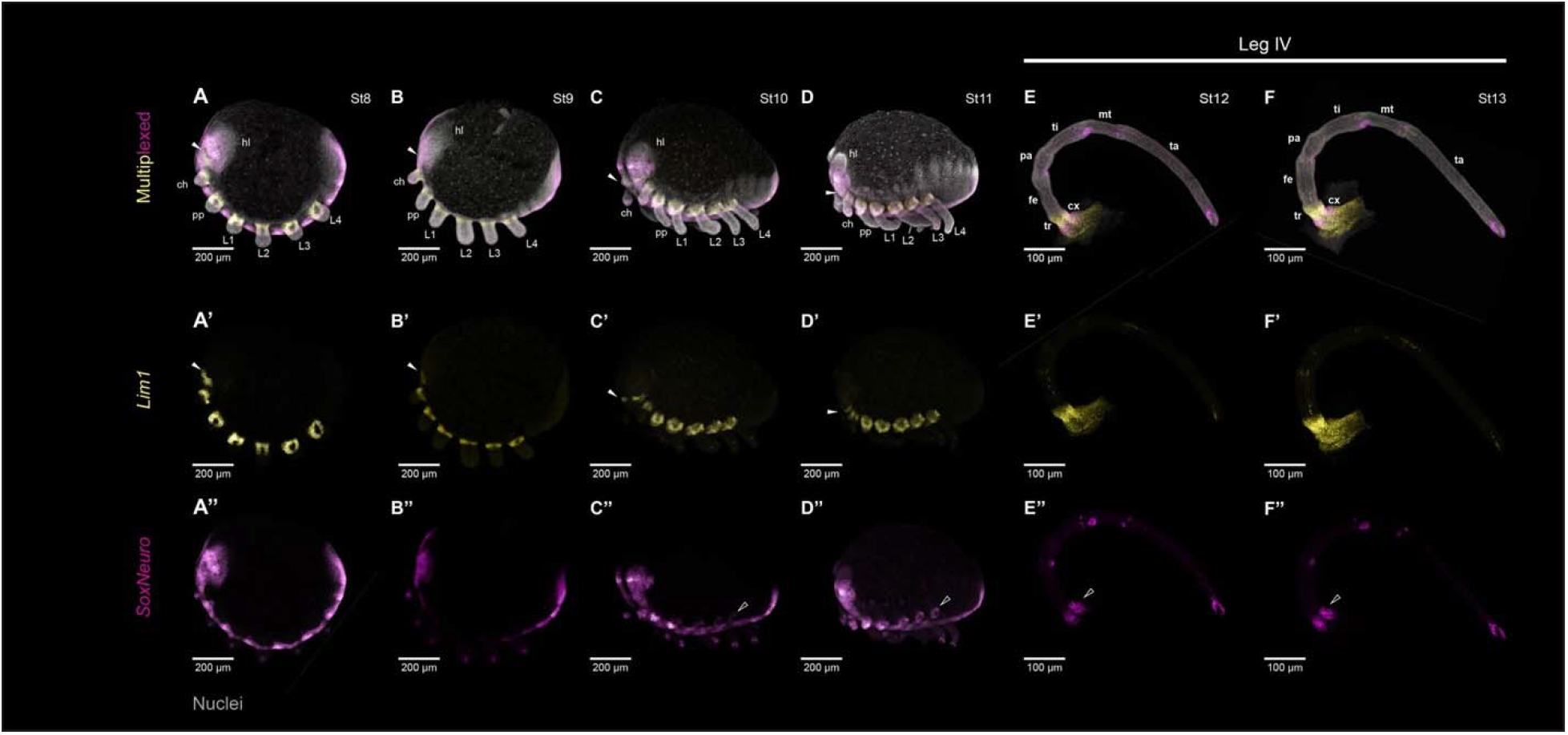
Po-Lim1 and Po-SoxNeuro are markers of proximal podomeres throughout P. opilio leg morphogenesis. (A-D) Nuclear counterstaining (grey) and multiplexed expression of *Po-Lim1* (yellow) and *Po-SoxNeuro* (magenta) in wildtype *P. opilio* embryos in successive stages of appendage development: stage 8 (A), stage 9 (B), stage 10 (C), stage 11 (D). (E-F) Dissected leg IV appendages of stage 12 (E) and stage 13 (F) embryos with multiplexed expression as in (A-D). Stages follow the nomenclature of (Gainett et al., 2022). (A’-F’) Isolated expression of *Po-Lim1*. (A’’-F’’) Isolated expression of *Po-SoxNeuro*. White arrowhead: non-appendicular *Po-Lim1* expression in the head lobes. Black arrowhead: proximal ring domain of *Po-SoxNeuro*. Abbreviations: hl – head lobe; ch – chelicera; pp – pedipalp; L1 – walking leg I; cx – coxa; tr – trochanter; fe – femur; pa – patella; ti – tibia; mt – metatarsus; ta – tarsus.

The proximal appendage expression of *Po-Lim1* remains largely static through later stages of appendage elongation with slight proximal expansion encompassing the margins of the lateral body wall (Fig. 2B-2D). By stages 12 and 13, *Po-Lim1* expression is heterogeneous, with the highest intensity of expression at the boundary of coxa and trochanter (Fig. 2E, 2F). Weaker patches of *Po-Lim1* expression are observed in minute distal domains of the appendages from stages 10 onward. These domains may correspond to morphogenesis of the internal elements of the appendage (e.g., muscle attachment sites; neural differentiation; Dolezal et al., 2024), in contrast to the broad domains throughout the appendage ectoderm that are associated with podomere formation or regionalization of the PD axis.

The expression of *Po-SoxNeuro* during appendage formation has been partially described in a prior work (Klementz et al., 2025). Briefly, *Po-SoxNeuro* exhibits high intensity expression in the ventral neuroectoderm in all surveyed stages. In the segmental appendages, *Po-SoxNeuro* first appears as a minute domain in the distal terminus in stage 8 embryos, maintained throughout development (Fig. 2A-2F). Proximal appendicular expression occurs by stage 10, appearing as a narrow ring domain at the distal boundary of the coxa that remains stable throughout appendage elongation toward stage 13 (Fig. 2C-F). As with *Po-Lim1, Po-SoxNeuro* additionally appears as small patches of expression in more distal territories. Expression in the distal terminus is comparable to domains observed in the head appendages of lineages such as myriapods, crustaceans, and onychophorans (Brenneis et al., 2021; Janssen et al., 2018; Pioro and Stollewerk, 2006). Likewise, distal tip expression and patchy domains in other appendage territories are characteristic of *SoxNeuro* expression in *D. melanogaster* (Crémazy et al., 2001).

### RNAi against Po-Lim1 incurs proximal leg defects, but no defects to the pretarsus

Consistent with prolonged localization to the proximal segments throughout appendage elongation, RNAi against *Po-Lim1* yielded deletions of the coxa and trochanter in both the pedipalps and walking legs (n = 91/295; 30.8%) (Fig. 3). The femur, tibia, and tarsus of both the palp and the leg exhibited truncations, with the femur appearing proximally adjacent to the body wall (Fig. 3B, 3D, 3F). No defects to the claws were observed in any RNAi-treated embryo (Fig. 3D, 3F).

**Fig. 3.**
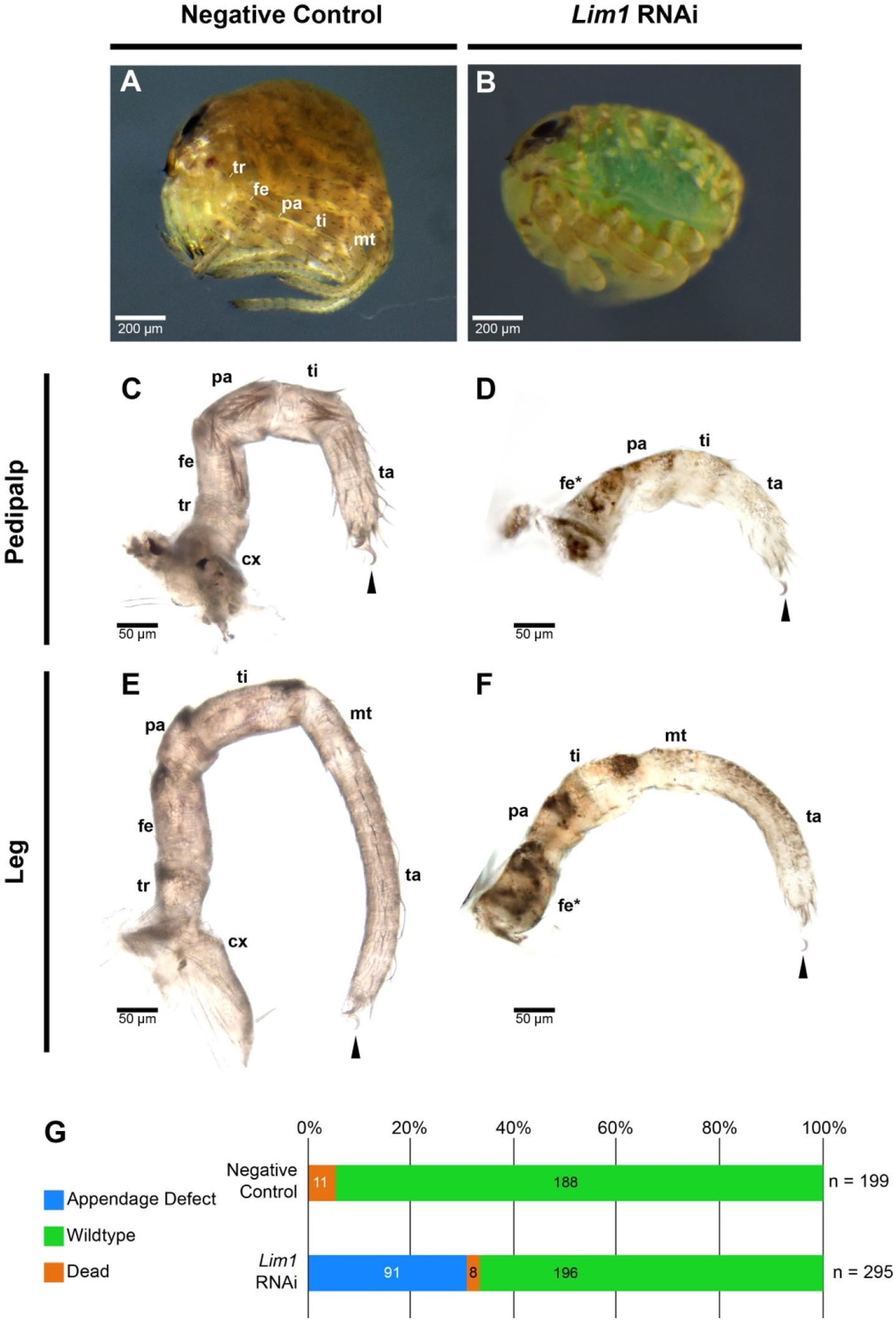
RNAi against Po-Lim1 results in deletion of proximal podomeres. (A) Negative control hatchling in lateral view. (B) *Po-Lim1* RNAi hatchling in lateral view. (C) Pedipalp of negative control hatchling. (D) Pedipalp of *Po-Lim1* RNAi hatchling. Note the absence of coxa and trochanter, with remnants of the body wall proximally adjacent to a truncated femur (asterisk). (E) Leg I of negative control hatchling. (F) Leg I of *Po-Lim1* RNAi hatchling. No defect to the tarsal claw was observed in any RNAi hatchling (black arrowhead). (G) Distribution of outcomes following *Po-Lim1* RNAi experiments. Abbreviations as in Fig. 2.

In order to validate the identity of proximally affected podomeres, with emphasis on the interpretation of the femur, we surveyed a subset of *Po-Lim1* RNAi embryos for the expression of *odd-skipped*, a well-characterized marker of developing joints between appendage segments (Heingård et al., 2019; Medina-Jiménez et al., 2025), and *SoxNeuro*, as a marker of the distal coxa (see Fig. 2C-F). Embryos were surveyed at stage 12, wherein podomere boundaries are discernible. Wild type embryos display a full complement of five or six ring domains of *Po-odd-skipped* corresponding to the boundaries between the six and seven segments of the pedipalp and walking leg, respectively (Fig. 4A-4C; Fig. S1). *Po-SoxNeuro* likewise exhibits overlapping expression with the *Po-odd-skipped* ring domain that demarcates the boundary between coxa and trochanter (Fig. 4A-C; Fig. S1). By contrast, the appendages of *Po-Lim1* RNAi embryos exhibit four ring domains in the pedipalp and five in the walking legs, as well as shortening of the total length of both appendage types, consistent with the interpretation of podomere deletion (compare Fig. 4D-F to Fig. 4A-4C). In the majority of cases, the ring domain at the proximal boundary of the femur directly abuts the body wall, consistent with a deletion of both coxa and trochanter (Fig. 4F; Fig. S1), while a minority of cases exhibit reduced remnants of the trochanter (Fig. 4E; Fig. S1). In addition, appendages of RNAi embryos do not exhibit proximal ring domains of *Po-SoxNeuro*, consistent with deletion of the coxa, rather than any medial or distal territory of the appendage.

**Fig. 4.**
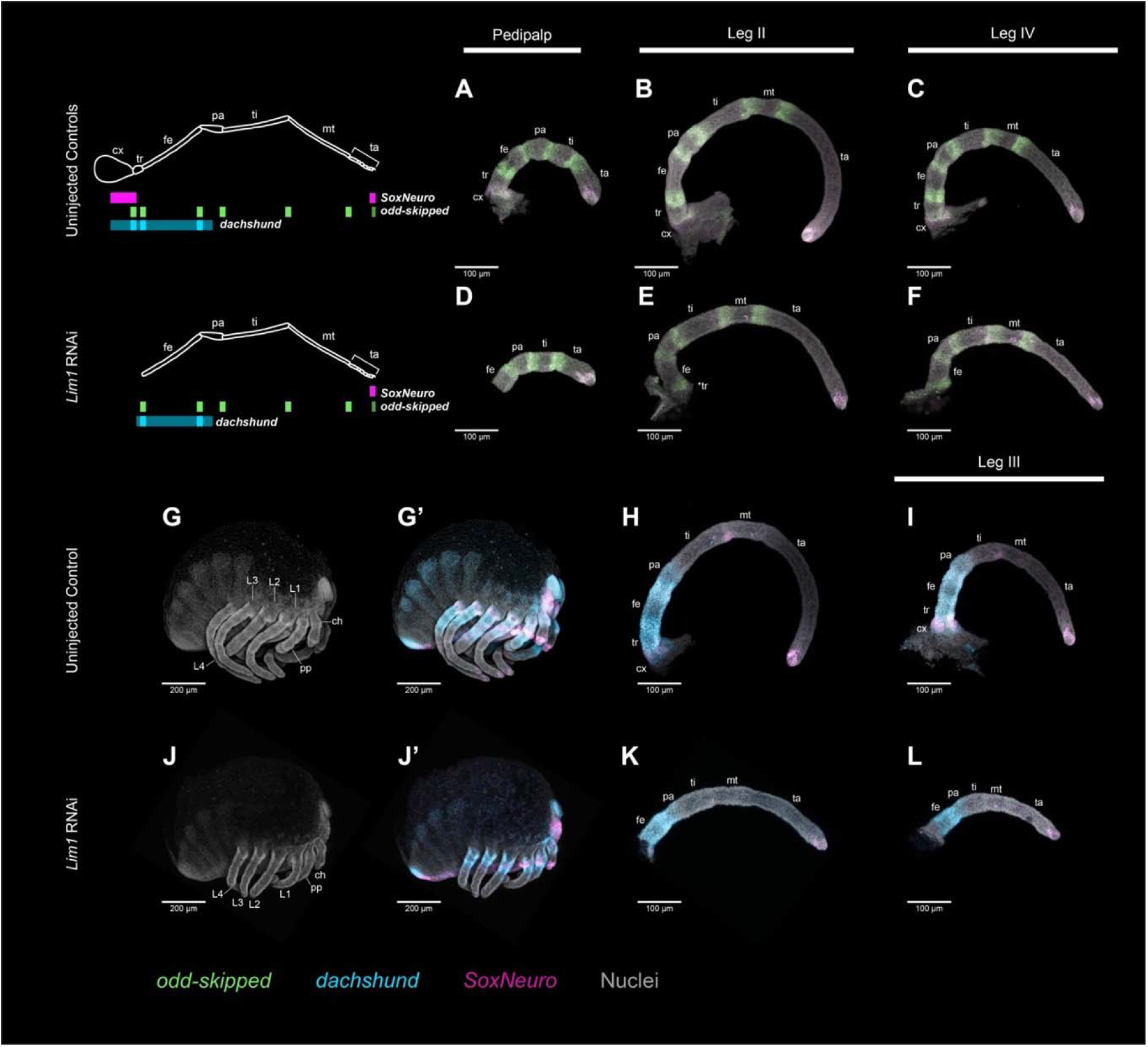
Disrupted expression domains of Po-SoxNeuro, Po-odd-skipped, and Po-dachshund in RNAi embryos validate proximal podomere deletions. Top left inset: schematic representation of gene expression differences between uninjected controls and *Po-Lim1* RNAi embryos. (A-C) Nuclear counterstaining (grey) and multiplexed expression of *Po-odd-skipped* (green) and *Po-SoxNeuro* (magenta) in the pedipalp (A), leg II (B), and leg IV (C) of a wildtype *P. opilio* embryo. (D-F) Multiplexed expression as in (A-C) in the pedipalp (D), leg I (E), and leg II (F) of a *Po-Lim1* RNAi embryo. (G) Isolated nuclear counterstaining in a stage 12 uninjected control embryo. (G’) Same embryo as in (G) with expression of *Po-dachshund* (cyan) and *Po-SoxNeuro*. (H-I) Leg II (H) and leg III (I) with expression of *Po-dachshund* and *Po-SoxNeuro* from an uninjected control embryo. (J) Isolated nuclear counterstaining in a stage 12 *Po-Lim1* RNAi embryo. (J’) Same embryo as in (J) with expression of *Po-dachshund* and *Po-SoxNeuro*. (K-L) Leg II (K) and leg III (L) with expression of *Po-dachshund* and *Po-SoxNeuro* from a *Po-Lim1* RNAi embryo. Abbreviations as in Fig. 2.

Next, a separate subset of *Po-Lim1* RNAi embryos was assayed for the expression of *dachshund*, a marker of the medial appendage territory during regionalization of the PD axis. While initially highly expressed throughout the medial appendage, by stage 12 *Po-dachshund* exhibits heterogeneous expression, with the highest intensities localized to the distal boundaries of the coxa, trochanter, and femur alongside intervening, weaker expression up through the proximal compartment of the patella (Fig. 4G-4I; Fig. S2). *Po-Lim1* RNAi yields both a truncation in the extent of *Po-dachshund* expression, as well as a decrease in the number of high-intensity expression domains (Fig. 4J-4L; Fig. S2). *Po-Lim1* RNAi embryos exhibit only two ring domains of *dachshund*, corresponding to the distal boundaries of the trochanter and femur; and no expression of *Po-SoxNeuro*, consistent with deletions of the coxa and proximal trochanter. These results are consistent with the morphology of the truncated podomere interpreted as a femur in *Po-Lim1* RNAi hatchlings (Fig. 3D, 3F).

### Lim1 *and* SoxNeuro *expression domains in the sea spider leg show a reversal of distal boundaries compared to the harvestman*

To assess whether the dynamics of *Lim1* and *SoxNeuro*, localized to and essential for the formation of coxa and trochanter, may be phylotypic of Chelicerata, we assessed the expression of single-copy homologs in the appendages of the sea spider *Pycnogonum litorale*, notable for the retention of three putative coxal homologs and no trochanter. *P. litorale* exhibits anamorphic development, initially hatching as a protonymphon larva with only three appendage pairs, the chelifores and the palpal and ovigeral larval limbs. Through successive molts, the walking legs appear alongside posterior body segments (Fig. S3). At instar II, the limb bud primordia of the first walking leg are still compressed beneath the cuticle (Fig. S3), but by instar III, the limb buds distinctly protrude from the body wall.

*Pl-Lim1* is expressed in a ring domain in the proximal territory of the outgrowing limb bud, with stronger intensity in the antero-dorsal portion of the ring (Fig. 5A, S3). This domain marginally overlaps with a distally abutting ring domain of *Pl-SoxNeuro*, resulting in a reversed relationship of the distal expression boundaries of both genes compared to the harvestman limb bud (Fig. 5A). Additionally, both *Pl-Lim1* and *Pl-SoxNeuro* are diffusely expressed at the distal limb bud tip. Proximal domains of *Pl-Lim1*, albeit weaker, are also observed in the proximal-most podomeres of the three anterior larval limb pairs, as well as strongly in the primordium of the second walking leg pair (Fig. 5A). However, *Pl-SoxNeuro* does not yet exhibit strongly localized expression in this territory (Fig. S3). In contrast to *Pl-SoxNeuro*, coexpression of *Pl-Lim1* in the proximal portion of the previously described *Pl-dachshund* territory (Klementz et al., 2024) supports conserved dynamics of these two genes with the harvestman through this early stage of appendage formation (Figs. 5C; S4)

**Fig. 5.**
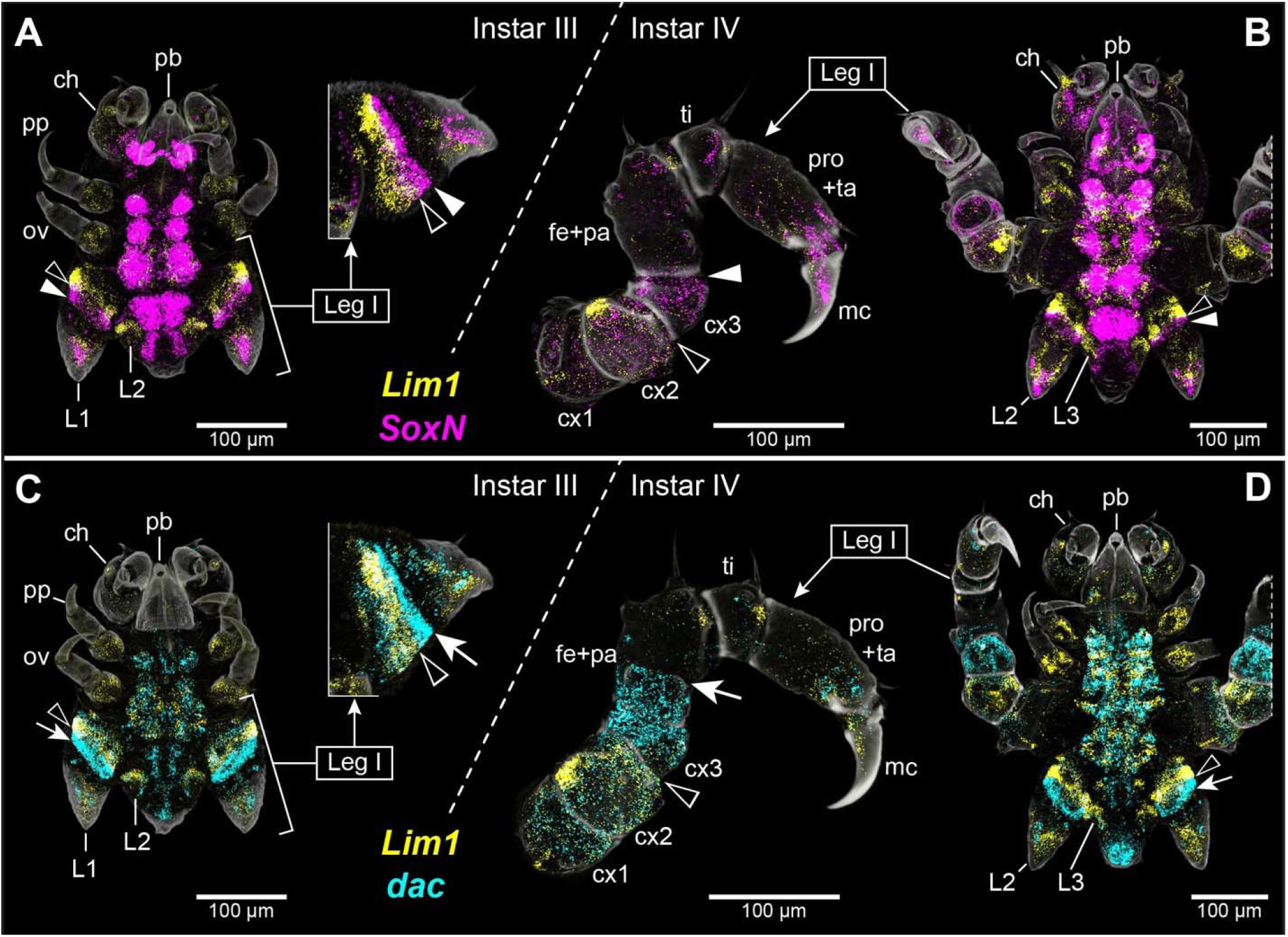
The expression dynamics of Lim1 and SoxNeuro homologs in the sea spider Pycnogonum litorale obfuscate alignment of proximal podomeres across Chelicerata. (A) Left: Instar III in ventral view with multiplexed expression of *Pl-Lim1* (yellow) and *Pl-SoxNeuro* (magenta). Right: magnification of leg I limb bud in lateral view. (B) Right: Instar IV in ventral view with expression as in (A). Left: magnification of leg I in lateral view. (C) Left: Instar III in ventral view with multiplexed expression of *Pl-Lim1* and *Pl-dachshund* (cyan). Right: magnification of leg I limb bud in lateral view. (D) Right: Instar IV in ventral view with expression as in (C). Left: magnification of leg I limb bud in lateral view. In all panels, cuticular autofluorescence is depicted in grey. The distal boundaries of *Pl-Lim1* (black arrowheads), *Pl-SoxNeuro* (white arrowheads), and *Pl-dachshund* (white arrows) expression domains are highlighted. Note that the rings of *Pl-Lim1* (black arrowheads) and *Pl-SoxNeuro* (white arrowheads) in instar III are much less obvious in early instar IV than those of *Pl-dachshund*. Abbreviations: pb – proboscis; ch – chelifore; ov – oviger; pro – propodus; mc – main claw; others as in Fig. 2.

By instar IV, the podomere boundaries are clearly discernible in the now functional first walking leg (Fig. 5B, 5D). In early instar IV specimens, the tissues in the recently molted leg still contain relatively few, spread-out cells, resulting in weaker expression signals compared to the compressed tissues in the limb buds (Figs. 5A, 5B; S3). *Pl-Lim1* expression remains strongly localized to the proximal podomere of the larval chelifore, palp, and oviger. In the walking legs, *Pl-Lim1* localizes most strongly to the second coxa, particularly the anterior-dorsal territory, with slightly more sparse expression through the remainder of the podomere (Fig. 5B, 5D). More limited expression patches are observed in the lateral margins of the first coxae as well as the distal margins of patella, tibia, and tarsus-propodus precursor, which may relate to muscles and/or their attachment sites at the podomere joints (Figs. 5B, 5D; S3; S4). Additionally, *Pl-Lim1* is expressed in the ventral compartment of the main claw. At the same stage, *Pl-SoxNeuro* maintains its strongest expression distally adjacent to *Pl-Lim1* in the proximal appendage, localizing to coxa 3 and the distal compartment of coxa 2. Early instar IV specimens show additional diffuse *Pl-SoxNeuro* expression in proximal coxa 2 and coxa 1, albeit with weaker intensity. With ongoing growth of the ectodermal leg tissues during instar IV, the *Pl-SoxNeuro* expression territories in the proximal leg become more distinct, including the ring domain in coxa 3 and distal coxa 2, as well as more localized expression patches in proximal coxa 2 and in coxa 1. Distal of the coxae, *Pl-SoxNeuro* is expressed in small patches in the femur-patella precursor, tibia, and tarsus-propodus precursor, and a broader domain in the main claw encompassing much of the medial and dorsal territories. As previously described, *Pl-dachshund* maintains its expression distal to coxa 2, extending up to the prospective boundary of femur and patella (Klementz et al., 2024) (Figs. 5D; S4)

### Expression of SoxNeuro homologs in a myriapod and crustacean exemplify evolutionary lability

Discrepancy in the expression domains of *Lim1* and, particularly, *SoxNeuro* between the harvestman and sea spider, could be suggestive of multiple competing hypotheses in the evolution of proximal appendage segmentation in chelicerate taxa. In the case of *Lim1*, a conserved role in the maintenance of both coxa and trochanter in an arachnid and several insect models, suggests that this function evolved in the arthropod common ancestor. Yet, for *SoxNeuro*, divergent expression domains, particularly in regard to the degree of co-expression with *Lim1*, could reflect either independent co-options of this gene in the appendage patterning of chelicerates, or alternatively developmental system drift in one or both lineages following the gain of appendicular expression domains in their common ancestor. The second hypothesis, however, is difficult to arbitrate given the dearth of *SoxNeuro* expression assays spanning the appropriate developmental time window and focusing on leg formation in mandibulate lineages. Surveys of *SoxNeuro* expression are currently limited to insects and a single exemplar of both myriapods and crustaceans, the pill millipede *Glomeris marginata* and the marbled crayfish *Procambarus virginalis*, respectively (Brenneis et al., 2021; Crémazy et al., 2001; Janssen et al., 2018). In insects, *SoxNeuro* is not expressed in the locomotory appendages. In *Tribolium, SoxNeuro* is expressed in the proximal territories of the mandibles and maxillae, similar to subsets of head appendages in the millipede and crayfish (Brenneis et al., 2021; Crémazy et al., 2001; Janssen et al., 2018; Pioro and Stollewerk, 2006). However, the prior works have not sampled stages of appendage development beyond limb bud, leaving the extent of *SoxNeuro* activity in the locomotory appendages of mandibulates unclear.

To supplement these prior surveys, we performed embryonic expression assays of *Lim1* and *SoxNeuro* in two additional mandibulates, the amphipod *Parhyale hawaiensis* and the centipede *Lithobius atkinsoni*. In the thoracopods of *P. hawaiensis, Ph-SoxNeuro* is expressed in a ring domain at the distal margin of the basis, the podomere immediately distal to the coxa (Fig. 6A). As in the sea spider, *Ph-SoxNeuro* is distally adjacent to a strong domain of *Ph-Lim1* in the coxa (Fig. 6B). Additional *Ph-SoxNeuro* expression occurs in scattered domains in the ischium, merus, and distal tips of propodus and dactylus, likely associated with appendicular mesoderm or sensory cells.

**Fig. 6.**
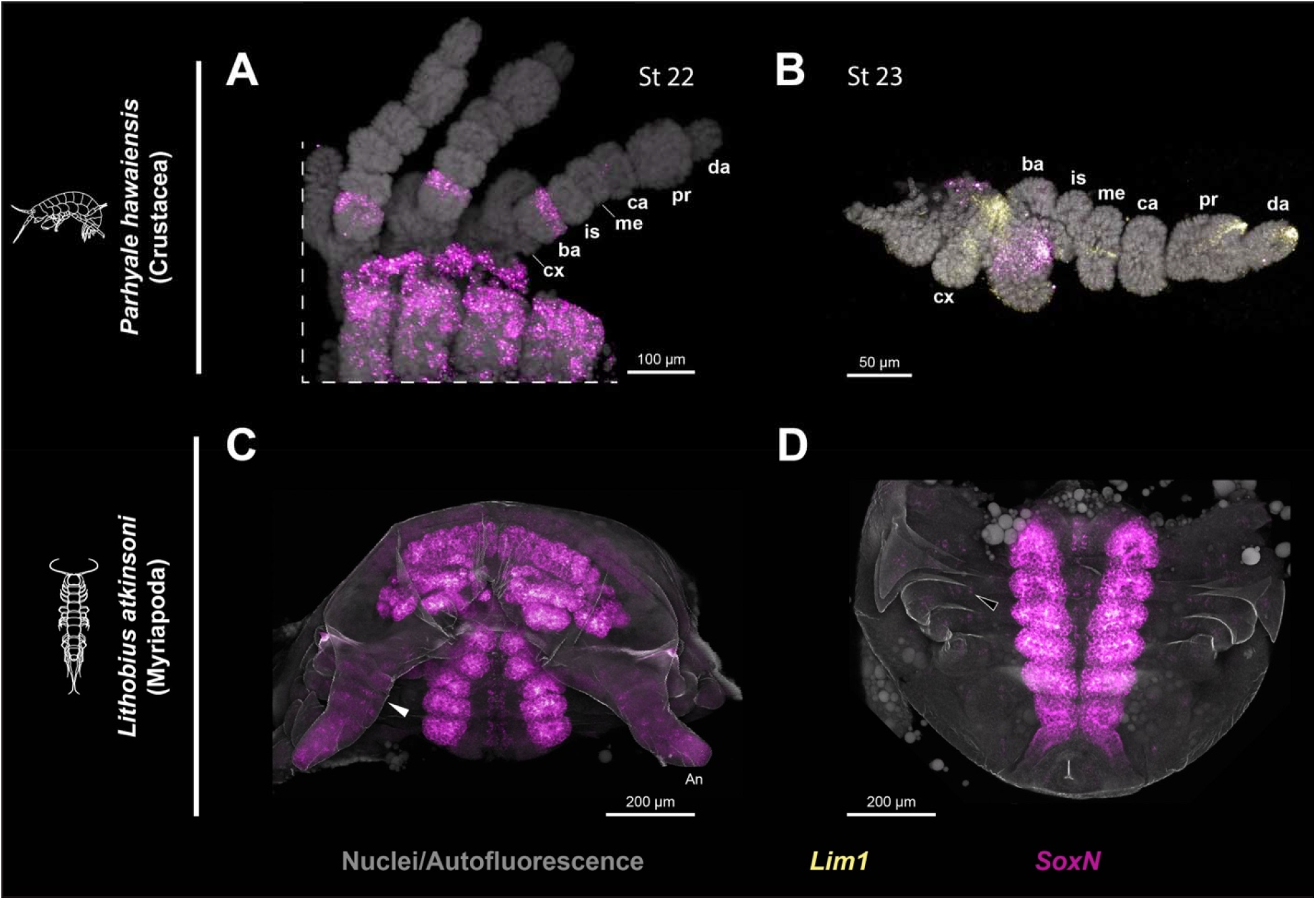
Mandibulate SoxNeuro homologs refute its utility as a comparative marker of proximal podomeres. (A) Expression of *Ph-SoxNeuro* (magenta) in a subset of thoracopods in a stage 22 embryo of the amphipod *P. hawaiensis*. (B) Co-expression of *Ph-Lim1* (yellow) and *Ph-SoxNeuro* in a stage 23 *P. hawaiensis* thoracopod. Note the localization of *Ph-SoxNeuro* to the basis, rather than coxa. Stages follow the nomenclature of (Browne et al., 2005) and nuclear counterstaining is depicted in grey. (C) Expression of *La-SoxNeuro* in the head and anterior trunk segments of the centipede *Lithobius atkinsoni*. (D) Detail of *La-SoxNeuro* in the posterior *L. atkinsoni* trunk segments. Note only minute, weak expression of *La-SoxNeuro* in the appendages. White arrowhead: putative ring domain of *Ls-SoxNeuro* in the antenna. Black arrowhead: example of putative sensory *La-SoxNeuro* expression. Grey coloration in (C,D) represents cuticular autofluorescence. Abbreviations: cx – coxa; ba – basis; is – ischium; me - merus; ca – carpus; pr – propodus; da – dactylus; an – antennae.

No homologs of *Lim1* were recovered in the developmental transcriptome of *L. atkinsoni*. In the appendages, strong *La-SoxNeuro* expression was recovered only in the antennae, with subtle ring domains proximally, grading into a strong cohesive domain toward the distal territory (Fig. 6C). In the walking legs, *La-SoxNeuro* exhibits only minute, weak spots of expression, perhaps indicative of developing sensory structures as in insect taxa (Fig. 6D).

### Expression dynamics of Irx2 challenge the presence of a cryptic precoxal homolog in chelicerates

A previous effort to homologize the leg segments of all four arthropod subphyla leveraged the transcription factors *Irx2* (*araucan*) and *pannier* to support inferences of an ancestral complement of eight leg podomeres including the terminal dactylus or pretarsi, in the appendages of all arthropods, with variable numbers of segments incorporated into the body wall in all lineages except sea spiders (Bruce, 2021). In this work, domains of *pannier* were inferred to demarcate the true body wall or medial tergum in all surveyed arthropods. In insects, three notable domains of *Irx2* expression occur distally of this *pannier* domain, with two of them proximal to the coxa, inferred to demarcate two precoxal podomeres incorporated into the body wall. By contrast, in crustaceans only one of the three domains localizes proximal to the coxa-body wall boundary, which was considered to indicate incorporation of only one precoxal podomere into the body wall of crustaceans. Similar dynamics of *Irx2* were observed in the tarantula *Acanthoscurria geniculata*, suggesting that chelicerates have likewise incorporated a cryptic precoxal podomere, inferences subsequently validated by surveys of *drumstick*, a marker of developing leg joints between true leg segments. These results were subsequently leveraged to support one-to-one alignment of podomeres across all arthropod lineages, including sea spiders (Bruce, 2021).

However, this hypothesis is predicated on the assumption that coxa 1 of sea spiders represents a homolog of the body wall-associated podomeres in arachnids and crustaceans, and the first precoxa of insects. In order to assess these assumptions, we surveyed both the harvestman and sea spider for the expression of *Irx2*. In the harvestman, *Po-Irx2* is exclusively expressed in the lateral body wall; no domains of expression are observed in the appendages (Fig. 7A-C’), contradicting interpretations of three conserved domains of *Irx2* across arthropod legs. Lateral expression adjacent to the legs, however, appears as two rough patches, which accords with interpretations of *Irx2* “bracketing” the putative precoxa of crustaceans and the tarantula. In the sea spider, however, *Pl-Irx2* is also restricted to the lateral body wall, expressed in a ring domain likely demarcating the lateral process, a tubular extension of the body wall upon which the leg is mounted (Fig. 7D-E). No domains of *Pl-Irx2* occur in the sea spider appendages, refuting the alignment of coxa 1 with a cryptic eighth leg podomere across extant lineages.

**Fig. 7.**
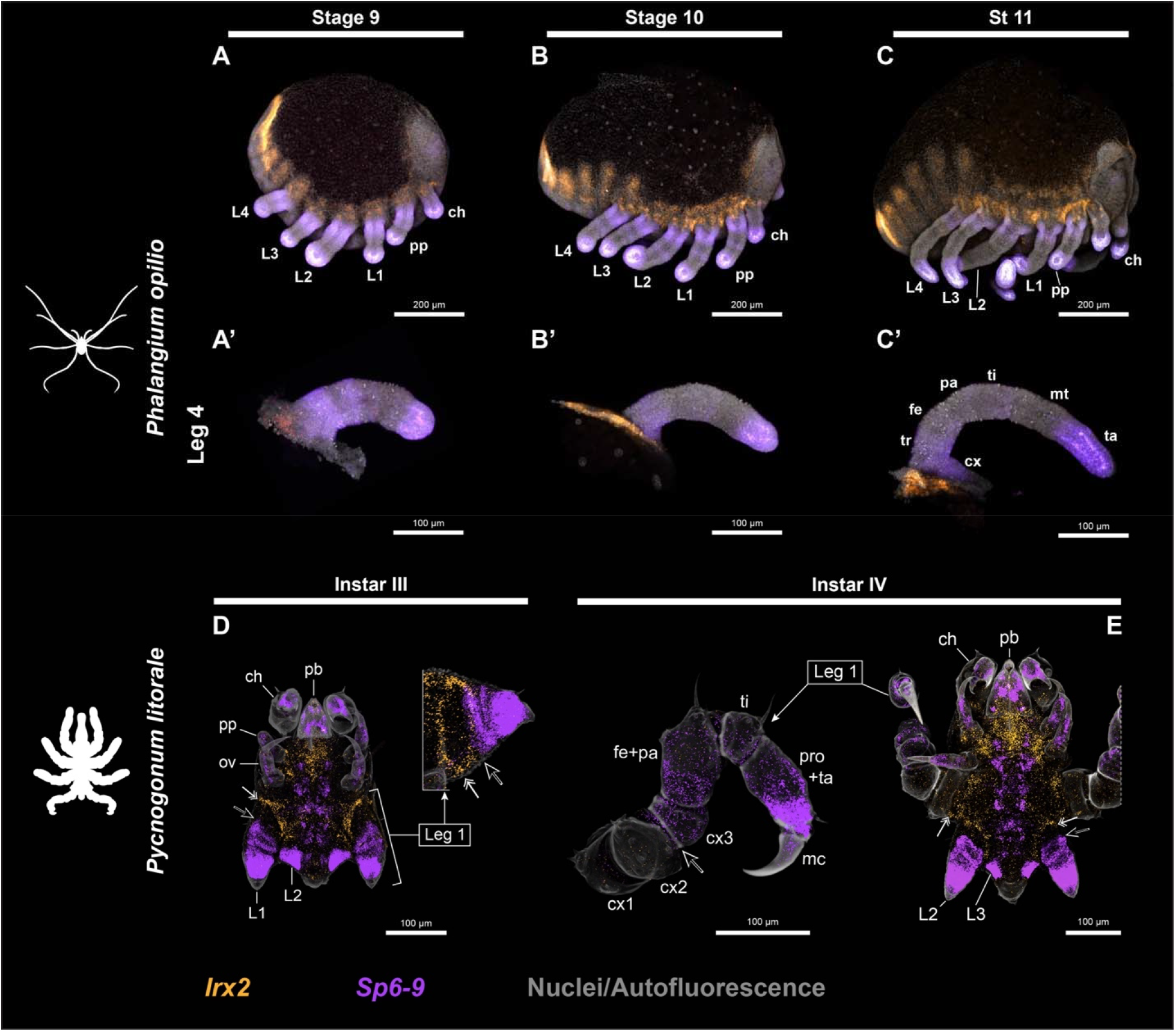
Comparative expression of *Irx2* and *Sp6-9* in the harvestman and sea spider suggests neither possess a cryptic precoxa but still confound proximal podomere alignment. (A-C) Multiplexed expression of *Po-Irx2* (orange) and *Po-Sp6-9* (violet) in wildtype *P. opilio* embryos in lateral view at stages 9 (A), 10 (B), and 11 (C). (A’-C’) Detail of *Po-Irx2* and *Po-Sp6-9* expression in the fourth walking leg of sampled stages. (D-E) Multiplexed expression of *Pl-Irx2* (orange) and *Pl-Sp6-9* (violet) in *Pycnogonum litorale* instars III (D) and IV (E). (D) Ventral view (left) with detail of leg I limb bud (right). (E) Ventral view (right) with detail of leg I (left). Note the proximal boundary of *Pl-Sp6-9* in the third coxa (black arrow), and most-distal *Pl-Irx2* expression in the non-appendicular lateral processes (dual white arrows). Abbreviations as in Figs. 2, 5.

### Proximal expression boundaries of Sp6-9 are not conserved across chelicerates

In a similar manner, the prior scheme suggested that conserved expression dynamics of *Sp6-9* across arthropod lineages supports homology of the six most distal leg elements, i.e., all podomeres distal to the second insect precoxa, distal to the crustacean and chelicerate coxa, and distal to the sea spider second coxa. As with *Irx2*, we endeavored to re-evaluate this claim via surveys of *Sp6-9* expression in the harvestman and sea spider leg. In the harvestman, *Po-Sp6-9* is expressed in multiple domains, including strongly in the tarsus, and faint rings at the distal boundaries of patella and tibia. Proximally, *Sp6-9* expression encompasses the coxa and trochanter, grading into progressively weaker expression in the proximal femur (Fig. 7A-C’). The spans of these domains conflict with the interpretations of Bruce (2021). Distally, the presence of *Po-Sp6-9* in the tarsus, but not the metatarsus, may conflict with the interpretation that both podomeres represent subdivided homologs of an ancestral, single tarsus podomere. Likewise, expression in the harvestman coxa, the seventh podomere in their reconstruction, contradicts the homology of the insect coxa, crustacean basis, and arachnid trochanter (Bruce, 2021). In the sea spider, *Pl-Sp6-9* exhibits similar expression dynamics (Figs. 7D-E, S5). In instar III, *Pl-Sp6-9* exhibits heterogeneous expression in the limb bud, with a strong distal domain throughout much of the limb, increasing to a more intense ring domain proximally. In the articulated leg of instar IV, expression is strongest in the distal tarsus-propodus precursor and the proximal main claw. Weaker expression also localizes to coxa 3 and femur, with marginally overlapping with the distal boundary of coxa 2. Sparse intervening expression between the proximal and distal domains also occurs. Notably, the proximal-most extent of the expression domains aligns with the interpretations of Bruce (2021), albeit with the same complication in the treatment of a singular unit composed of both tarsus and propodus.

## Discussion

In recent years, our understanding of the genetic mechanisms that underlie the evolution of chelicerate appendage patterning has steadily advanced through the lens of the model harvestman, *Phalangium opilio*. Functional datasets and broad comparative expression surveys drawn from this system now underscore both conserved ancestral and derived features of chelicerate appendage development (Gainett et al., 2021; Klementz et al., 2024; Medina-Jiménez et al., 2026; Sharma et al., 2013). However, only limited data are available on the patterning of proximal territories in chelicerates (Klementz et al., 2024; Sharma et al., 2015). Here, we investigated proximal appendage patterning in chelicerates using a combination of functional and comparative data and brought these data to bear on our understanding of arthropod leg evolution.

### A partly conserved role for Lim1 in coxal patterning in an arachnid

RNAi against *P. opilio Lim1* consistently resulted in deletions of both the coxa and trochanter, the two most proximal podomeres of the pedipalps and walking legs. This result is partly consistent with phenotypic outcomes following *Lim1* depletion in insect models, particularly in the proximal territory of the appendages. In both *Drosophila* and *Tribolium*, loss of function experiments consistently yield deletions of the coxa and/or deletions of podomeric boundaries between coxa-trochanter and trochanter-femur (Angelini et al., 2012; Tsuji et al., 2000). In contrast to the harvestman, however, no deletions of the trochanter are apparent following *Lim1* depletion in insects, consistent with a lack of expression of *Lim1* in the insect trochanter. This may represent an instance of developmental system drift, wherein the underlying genetic mechanisms responsible for patterning homologous structures (in this case, the trochanter) diverge over evolutionary time. This outcome echoes a prior case study demonstrating divergent patterning of the chelicerate tarsus (Klementz et al., 2026). Nonetheless, both insect and arachnid *Lim1* are essential for the establishment of the proximal-most podomere, suggestive of a conserved, ancestral role across Arthropoda.

Outside of proximal segments, however, insects often exhibit aberrant morphology of the distal femur and proximal tibia following *Lim1* disruption, consistently appearing with a bent appearance (Angelini et al., 2012). These defects maintain segmental boundaries between femur and tibia, an unlikely result if *Lim1* were necessary for the specification of this territory. Rather, this defect has been attributed to a distinct role for *Lim1* in the development of leg segment musculature. Cell lineage tracing experiments in *Drosophila* have demonstrated that in late larval stages, mesodermal precursor cell populations give rise to muscles in distinct territories of the walking legs (Guillermin et al., 2026). Mesodermal precursors at the center of the leg imaginal disc produce muscles of the femur and tibia, whereas peripheral cell populations give rise to muscles of the coxa, trochanter, and body wall. Notably, these differences are attributable to divergent complements of transcription factors expressed in the two populations; central cells express transcription factors such as *bifid* and *Lim1*, while the peripheral express *unplugged*. Further perturbations show a differential impact of *Lim1* primarily on the morphology of femur, while *bifid* affects the morphology of the tibia, explaining the more severe defects to the femur.

In the harvestman, disruption to the femur is subtle, appearing as a proximal truncation, as evidenced by reduced *dachshund* expression domains. Yet in all stages examined, none appeared with the pronounced bend characteristic of insect *Lim1* loss-of-function phenotypes. *Lim1* expression in the femur appears to be restricted to internal mesoderm (see Fig. 2F), rather than the ectodermal expression characteristic of transcription factors involved in the patterning of distinct territories. However, it cannot be ruled out that *Lim1* may have a role in specifying the proximal femur early in limb development, wherein podomere boundaries cannot be clearly delimited and thus the extent of *Lim1* expression is unclear. It is also plausible that this expression in the femur mesoderm may represent an analogous function to the specification of femoral muscle, as in insects. Differences in the severity of femur deformation may be the result of the divergent mechanisms of appendicular muscle development between chelicerates and insects. In *Drosophila*, mesodermal precursors acquire distinct muscular identities in late stages of larval development, prior to fusion into muscle fibers, without regard to patterns established by tendons or other internal appendicular tissues, exemplified by misplaced tendon insertions following *bifid* depletion (Guillermin et al., 2026). This would explain the pronounced defects following pupal *Lim1* RNAi in *Tribolium*, the stage at which distinct muscle identities are ascribed to the mesodermal precursors (Angelini et al., 2012), but extrapolating this to chelicerates is challenging, as myogenesis has not been formally characterized in any chelicerate to date. Prior works suggest that appendicular musculature is derived from coelomic extensions into the appendages from paired somitic pouches along the body segments (Moritz, 1957). These mesodermal extensions are visible in *Phalangium* limb buds upon outgrowth from the body wall (Gainett et al., 2022). This may represent a similarity to the migration of mesodermal precursors into the appendages of vertebrates, wherein locally secreted morphogens and established tissues determine their positioning and ultimate fate (Hasson et al., 2010). Thus, as our *Lim1* experiments targeted stages just prior to the establishment of limb buds, we may have disrupted positional information provided by *Lim1* for only a small window of time, such that recovered expression may have rescued any further defects of the internal mesoderm as the leg elongates. Testing this scenario in future will require a detailed characterization of chelicerate myogenesis.

### Evolution of Lim1 function in claw patterning may reflect yet another case of developmental system drift

In the fruit fly, the pretarsus and its claws require the concerted action of *Lim1, aristaless*, and *clawless* for normal development (Kojima, 2017; Pueyo et al., 2000). Early in the establishment of the leg imaginal disc, *aristaless* and *clawless* are expressed in the presumptive pretarsus, with *Bar* expressed in the presumptive distal tarsus. These distal, *aristaless-* and *clawless-*positive cells activate the expression of *Lim1* which represses *Bar*, which in turn represses *aristaless* and *clawless*. This reciprocal repression, mediated by *Lim1*, allows for the precise separation of tarsus and pretarsus (Pueyo et al., 2000; Tsuji et al., 2000). We have previously demonstrated, however, that *clawless* is not required for the specification of the harvestman pretarsus and its claws (Klementz et al., 2026). Instead, RNAi-mediated knockdowns yield disruptions to tarsal patterning. This functional divergence is associated with divergent expression dynamics. While *clawless* is broadly expressed throughout the distal appendage early in limb bud formation, it localizes strongly to the boundary of metatarsus and tarsus at later stages, no longer co-expressed with *aristaless* in the distal terminus. Broader comparative surveys of *clawless* expression across chelicerates highlighted this tarsal-specific function as ancestral to the subphylum.

Part of our motivation for investigating the function of *Lim1* in a chelicerate was to assess the evolution of the distal leg patterning program across Arthropoda (Setton and Sharma 2018; Klementz et al. 2024). Yet, as with *clawless*, RNAi against *Lim1* in the harvestman did not yield any defects of the distal claws. This suggests that the developmental genetic mechanisms underlying distal appendage formation have diverged between mandibulates and arachnids. This is supported further by expression surveys of *Bar*; harvestman *Bar* is expressed in the distal appendage terminus alongside *aristaless*, suggesting divergent mechanisms of separation of tarsus and pretarsus as well (Klementz et al., 2025). Curiously, *Lim1* RNAi also fails to produce defects to the pretarsus or claws in the flour beetle *Tribolium castaneum* (Angelini et al., 2012). Without broader sampling of insect lineages, it cannot be ruled out that the dynamics of *Lim1* in *Drosophila* may reflect a lineage-specific recruitment toward distal appendage patterning.

In the harvestman, one of the few remaining candidate genes in the claw-patterning network known from the *Drosophila* literature is *aristaless*. It is possible that chelicerate *aristaless* may have subsumed the role of the three *Drosophila* transcription factors, but no functional data exist to date for any chelicerate *aristaless* homolog.

### Functional dynamics of Lim1 do not support a simple one-to-one alignment of podomeres

Developmental system drift is broadly correlated with large phylogenetic distances for many developmental processes. However, it can obscure the underlying homology of morphological structures if the corresponding patterning mechanisms diverge beyond recognition. In the present case, conserved dynamics of many of the genes responsible for the initial regionalization of the PD axis are suggestive of a conserved, core gene regulatory network responsible for the formation of arthropod appendages. Conservation of these patterns has inspired a recent hypothesis for the alignment of podomeres across all arthropod lineages, based upon comparative gene expression assays of known PD axis-patterning transcription factors (Bruce 2021). Using a combination of the expression domains of six genes, Bruce (2021) proposed a one-to-one homology of all arthropod podomeres and deduced an ancestral complement of eight total podomeres, a condition considered to be retained in the walking legs of extant sea spiders. To accommodate both the expression patterns and unequal numbers of podomeres in extant arthropods, this alignment scheme invokes the incorporation of one to two podomeres into the body wall of all non-sea spider lineages. A precoxa is presumed to be present in arachnids, myriapods, and crustaceans, and two precoxae in the legs of insects. One drawback of this model, as previously discussed by us (Klementz et al. 2024), is that some genes do not accord with the proposed alignment; *dachshund* in particular exhibits demonstrable divergence of expression and function between the chelicerate and insect leg, regardless of the inferred podomere alignment scheme.

The results of the present study further question the underlying logic of the proposed alignment of Bruce (2021) and the inbuilt assumptions about the fidelity of expression patterns to specific podomeres. For example, under the newly proposed alignment scheme, the insect coxa and the arachnid trochanter are one pair of homologs; the insect trochanter is the presumed equivalent of the arachnid femur; and the insect femur is the equivalent of the arachnid patella. If so, depletion of *Lim1* (which causes loss of the insect coxa and severe truncation of the insect femur) should have yielded loss of the arachnid trochanter, severe truncation of the arachnid patella, but no effect on the arachnid coxa or femur. Instead, we observed that both the coxae and the femora are similarly affected by the loss-of-function experiments in both insects and the arachnid. The proposed alignment would also have predicted a deletion of the second insect precoxa in *Lim1* mutants, yet no defects to the lateral body wall were reported in either *D. melanogaster* or *T. castaneum* experiments. The insect experiments also yielded defects to the proximal tibia, an outcome not observed in any harvestman RNAi embryo. The partial overlap between insect and arachnid *Lim1* functional data thus disfavors the out of register alignment hypothesis of Bruce (2021). These new data are also congruent with previous works showing that the arachnid patella is a novel, lineage-specific gain of a podomere that is restricted to chelicerates, preserving the traditional alignment of the femora and tibiae of insects and arachnids (Turetzek et al. 2016; Klementz et al. 2024).

### Broader sampling of arthropod taxa further disfavors a simple one-to-one alignment of podomeres

A second drawback of the Bruce (2021) model is that, while comparative expression and functional data were available for some lineages of arthropods, no expression data were available for any sea spider to substantiate inferences of segment alignments and presumed range of expression patterns. In the past few years, advances in functional and genetic tools have broken this impasse and facilitated examination of gene expression in the sea spider model *Pycnogonum litorale* (Klementz et al. 2024, 2025; Papadopoulos et al. 2025). We brought this system to bear upon the question of leg evolution and tested the predictions of the Bruce (2021) model (Fig. 1C, Fig. 8A).

**Fig. 8.**
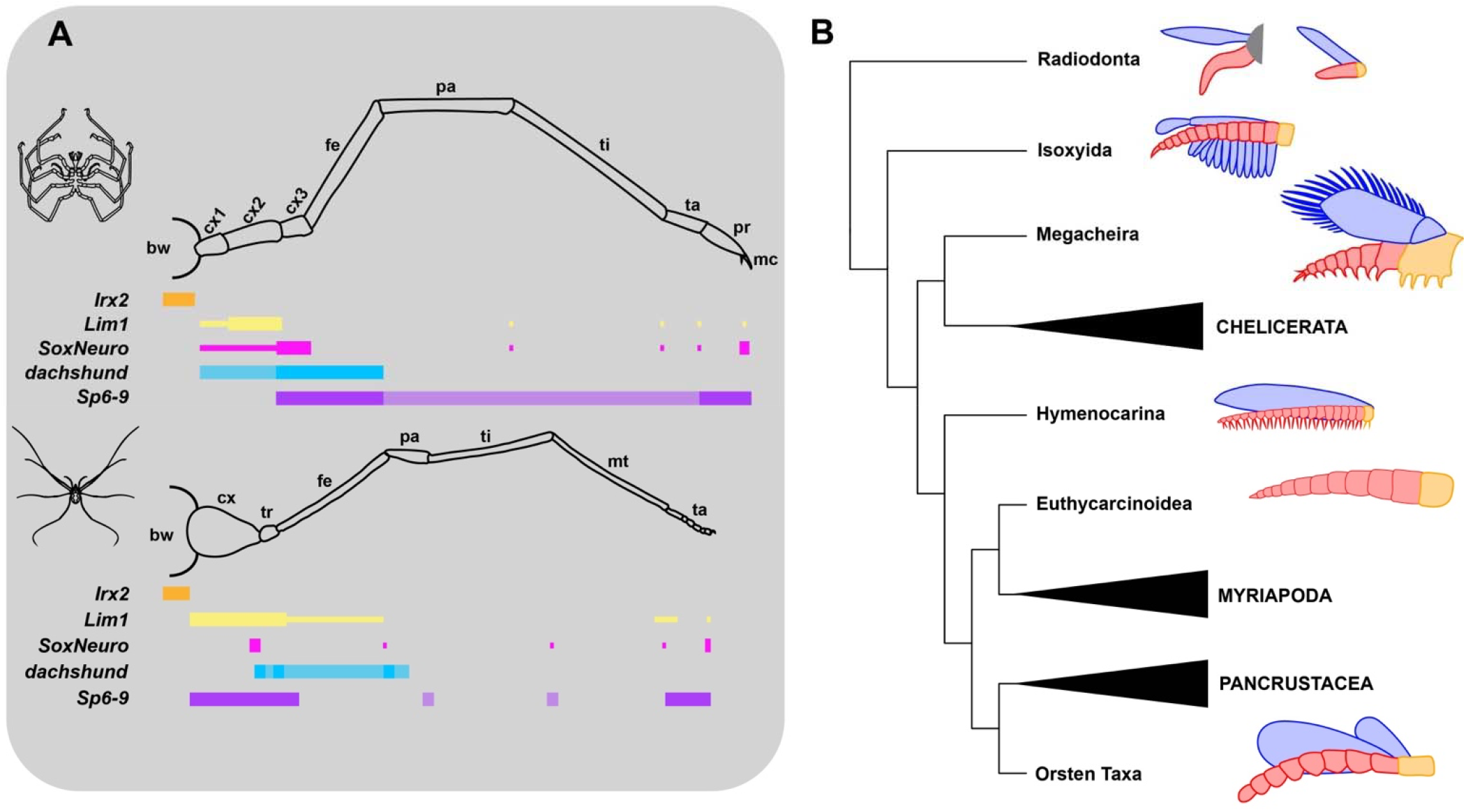
Homoplastic patterns of gene expression and inflated podomere counts in fossil taxa refute simplistic reconstructions of arthropod appendage evolution. (A) Schematic representation of PD axis-patterning transcription factor expression in the sea spider (top row) and harvestman (bottom row) walking leg. Thin boxes correspond to expression domains associated with only a subset of podomere tissue (e.g., mesoderm, muscle attachment sites). Abbreviations as in Fig. 2. (B) Simplified arthropod phylogeny including notable stem arthropods (Radiodonta, Isoxyida) and stem group representatives of all extant subphyla. Phylogeny modified from (Ortega-Hernández, 2026). Simplified diagrams of trunk appendages are provided to the right of stem-group taxa with endopod depicted in red, exo- and epipods in blue, and basal segment in yellow. Note that each appendage bears more than the eight podomeres presumed to be ancestral to all arthropods (Bruce 2021). Isoxyida appendage modeled after *Isoxys curvirostratus* (Zhang et al., 2021). Megacheiran appendage modeled after *Leanchoilia persephone* (García-Bellido and Collins, 2007). Hymenocarine appendage modeled after *Odaraia alata* (Alejandro Izquierdo-López and Jean-Bernard Caron, 2024). Euthycarcinoid appendage modeled after *Heterocrania rhyniensis* (Anderson and Trewin, 2003). Stem-pancrustacean Orsten taxa appendage modeled after *Ercaicunia multinodosa* (Zhai et al., 2019).

*Lim1* expression in the sea spider further challenges the simple one-to-one podomere homology scheme. In *P. litorale*, the strong antero-dorsal expression patch at the distal boundary of the *Lim1* ring domain of the limb bud can be safely traced to the dorso-anterior coxa 2 in the functional leg of instar IV. A complete ring-shaped *Lim1* domain is no longer readily discernible at this stage, but lower *Lim1* signal is nonetheless also retained in the remaining coxa 2 as well as coxa 1. While strong *Lim1* expression in the two most proximal podomeres during sea spider walking leg elongation and differentiation is shared with the harvestman, it does not conform with the *Lim1* domain expected for the podomere homology of Bruce (2021), which would predict its distal boundary in coxa 3 instead of coxa 2.

The limitation of strong *Lim1* expression from a continuous ring domain in the limb bud to an antero-dorsal patch in coxa 2 of the functional sea spider leg questions a continued PD appendage axis patterning function for *Lim1* after the proximal podomeres have been fully formed. To assess whether this local restriction of *Lim1* expression is specific to sea spiders, comparative data for later stages of leg differentiation in other arthropod lineages will be necessary. Given its persistent expression in the ectoderm, however, an exclusive role of *Pl-Lim1* in the development of musculature in coxa 2 seems unlikely, as underscored by the fact that functional intrinsic muscles sets of the proximal podomeres are already differentiated by this stage (Brenneis et al. 2023).

We had previously suggested that *SoxNeuro* could serve as a potential marker of proximal podomere identity, based on ring domains present in the distal harvestman coxa and sea spider coxa 3 (Klementz et al., 2025). However, differences between the two lineages in regard to co-expression of *Lim1* challenge its utility in assigning podomere homologies. This inspired us to test the evolutionary dynamics of *SoxNeuro* expression across the major lineages of Arthropoda, with emphasis on mandibulates. We found that the centipede lacks expression in the appendages outside of minute domains putatively corresponding to developing ectodermal sensory structures, according with prior expression surveys in a millipede and insect taxa (Crémazy et al., 2000; Luo et al., 2026; Pioro and Stollewerk, 2006; Rizzo and Bejsovec, 2017). In the amphipod, however, *SoxNeuro* is expressed in a ring domain along thoracopods, but at the distal boundary of the basis. This localization may be shared with crayfish, for which *SoxNeuro* expression was reported at the bifurcation point of exo- and endopod in early thoracopod stages, but confirmation in more advanced stages with defined podomere borders is currently lacking (Brenneis et al. 2021). Notably, according to the homology scheme proposed by Bruce (2021), the distal expression boundaries of *SoxNeuro* and *Lim1* in amphipod and sea spider legs coincide in suggested podomere homologs (crustacean coxa and sea spider coxa 2; crustacean basis and sea spider coxa 3). However, this similarity is neither shared with the harvestman, nor with myriapods (see above). In conjunction with the currently inconclusive presence of *SoxNeuro* in the appendages of a spider (Bonatto Paese et al., 2018), the available data suggest that *SoxNeuro* expression is inconsistent or labile across arthropods, with likely independent gains of appendicular ring domains in the amphipod, sea spider, and harvestman.

Another of the genes crucial to the homology scheme of Bruce (2021) is *Sp6-9*, used to support homology of the six distal-most podomeres across lineages. While the expression of *Sp6-9* is heterogeneous in its domain across taxa, the strong proximal and distal domains serve to bracket this appendage territory, which is deleted upon knockdown in spiders (Königsmann et al., 2017). We have subsequently reproduced this domain in our previous works in schematic representations of arthropod transcription factor expression. Here, we reexamined this proposition alongside comparative expression of the sea spider. The span of *Sp6-9* expression in the sea spider accords with Bruce (2021), with expression detected from coxa 3 through the main claw. But in the harvestman, *Sp6-9* is in fact expressed proximally through the coxa, not the trochanter as previously depicted; there is a mismatch between the span of the expression pattern and the functional domain of this gene. This further complicates assignments of homology of these proximal podomeres.

Despite the mismatches described above, the basis of Bruce (2021)’s scheme is the hypothesis of eight ancestral podomeres of the arthropod leg. Bruce (2021) supported the claim of internalized podomeres in non-sea spider arthropods via surveys of both *pannier*, a marker of the true body wall, and *Irx2* (*araucan*); *Irx2* expression distal to the *pannier* domain, but proximal to the appendages themselves, is perceived to demarcate the podomeres incorporated into the body wall of all non-sea spider arthropods. This idea is itself a derivation of an older hypothesis postulating the presence of a precoxa in *Tribolium* (Coulcher et al., 2015). This alignment predicts comparable expression of *Irx2* in the sea spider coxa 1 and in the putative arachnid precoxa; smaller domains of expression are expected in the next two distal podomeres across arthropods. While the harvestman does express *Irx2* in the body wall, it lacks the more distal domains of expression in coxa and trochanter predicted by Bruce (2021). Part of this confusion may stem from the use of a tarantula as a chelicerate exemplar in Bruce (2021). As members of Arachnopulmonata, spiders bear an ancestral whole genome duplication, facilitating additional copies of many genes (Schwager et al., 2017). Bruce (2021), however, survey the expression of only one *Irx2* gene, yet other spider lineages bear two copies, including the araneomorph *Parasteatoda tepidariorum* and another tarantula *Aphonopelma hentzi* (Setton et al., 2024). It is thus unclear which *Irx2* copy was surveyed, a particular concern given that the presence of multiple gene copies may relax selection pressure allowing for the accelerated evolution of one of the two copies, which may produce divergent expression domains associated with potential new functions (Gout et al., 2023; Holland et al., 2017). But more problematic is the complete absence of *Irx2* in any territory of the sea spider appendage and its restriction to the body wall (the same pattern as the harvestman). This fundamentally challenges the inference that the coxa 1 of sea spiders has been incorporated into the body wall of all other arthropods.

### The fossil record does not support an eight-podomere limb in the common ancestor of extant Arthropoda

The totality of these data suggests that subsets of PD axis patterning genes produce conflicting interpretations of segmental affinity across lineages, a consequence of a high degree of lability in their expression domains (Fig. 8A). Such lability had already been well characterized in comparisons of chelicerates and mandibulates (Klementz et al., 2026, 2025b, 2024; Sharma et al., 2015; Turchyn et al., 2011; Turetzek et al., 2017). A simpler— and more commonly accepted—thesis is that gene expression domains evolve like any other trait and sometimes cannot be used to establish simple homology schemes between structures, particularly within the arthropod PD axis. Attempting to force expression patterns and sparse functional datasets into reductive scenarios in the absence of external validation incurs implausible outcomes. In this case, an eight-podomere ancestral leg model effectively implies that structures like the coxa, trochanter, and femur have all evolved independently—including their specific arrangement of joint types and articulation—in chelicerates, myriapods, and insects. An over-reliance on expression patterns also hazards a potential circularity; expression domains are used to assign homologous podomeres, those assignments are then used to argue that the expression domains are evolutionarily conserved, and the resulting correspondence becomes evidence for the reconstructed ancestral appendage. Unless independent anatomical or phylogenetic criteria support the assignments, the inference risks becoming self-reinforcing.

In the present case, the use of developmental genetic data to reconstruct an ancestral, eight-segmented leg is inconsistent with the fossil record (Fig. 8B). Within the chelicerate stem group, the megacheiran genus *Leanchoilia* is described as bearing trunk endopods of nine podomeres plus a terminal claw (García-Bellido and Collins, 2007). Within the crown group, stem groups of extant orders as well as extinct orders bear a higher complement of podomeres. The sea spider stem group representative *Palaeopantopus maucheri* bears at least ten podomeres in its walking legs II-IV, with ambiguous additional segmentation of the distal appendage territory (Sabroux et al., 2024). Legs II-IV of other sea spider stem group fossils are subdivided into nine podomeres plus the main claw (Siveter et al. 2023; Sabroux et al. 2024). Eurypterida exhibit variation in the number of podomeres, ranging from seven to nine, with higher counts associated with prosomal appendages V and VI (Bicknell et al., 2020; Lamsdell, 2025; Lamsdell et al., 2015).

The mandibulate side of the tree is no different in this regard. Hymenocarines, putatively a stem group mandibulate lineage, often bear endopods with exaggerated podomere counts with 14 in *Balhuticaris voltae* (Izquierdo-López and Caron, 2022), 20 in *Pakucaris apatis* (Izquierdo-López and Caron, 2021), and 20 in *Odaraia alata* (Izquierdo-López and Caron, 2024). Similarly, the stem group lineage of Myriapoda, Euthycarcinoidea, was recently identified through exceptional preservation of the head capsule (Edgecombe et al., 2020). One exemplar, *Heterocrania rhyniensis*, bears trunk appendages consisting of 16 podomeres (Anderson and Trewin, 2003). Fuxianhuiids, alternatively considered stem or crown group mandibulates with close affinities to myriapods (Strausfeld et al., 2026, 2025), are often described with appendages consisting of more than a dozen podomeres (e.g., Yang et al., 2018). Putative stem and crown group pancrustaceans, such as *Waptia fieldensis* and *Yicaris dianensis*, respectively, bear more than eight podomeres in trunk appendages (Vannier et al., 2018; Zhang et al., 2007). An integrative view of paleontological data with extant character states therefore supports a model of independent reductions of podomeres in chelicerates and mandibulates.

A broader problem with a reductive, one-to-one segment alignment scenario is that it demands implausible reconstructions of limb evolutionary history. For example, an out of register alignment of coxae across the phylum requires the corollary that exopods cannot be homologous across Arthropoda. Bruce (2021) proposed that exopods are associated with the sixth podomere of the crustacean appendage, but the eighth podomere of fossil chelicerates such as *Offacolus* and *Dibasterium* (the equivalent of a putative precoxa in modern horseshoe crabs). The biramous arthropod leg must therefore have evolved at least twice. This non-parsimonious reconstruction draws no support from the fossil record, which substantiates a single gain of the biramous condition in the common ancestor of Deuteropoda, followed by the formation of podomeres along the PD axis (Boxshall, 2004; Fu et al., 2022).

The possibility that chelicerates and mandibulates have undergone independent reductions in their respective podomere complement explains well the discrepancies between gene expression patterns across the phylogeny, particularly in proximal podomere patterning. The evolutionary transformation series of arthrodized appendages suggests that the ancestral appendage of Deuteropoda comprised a large number of homonomous podomeres (e.g., in pectocaridids), in addition to exopods and epipods (Fu et al. 2021). The significance of the secondary branches is that they provide positional information for understanding the sequential reduction of podomeres (i.e., the origin of podomeres like the coxa). The inference of podomere reduction in crown-ward clades of deuteropods, both in the proximal (exopod- or epipod-bearing) and the distal territory, is therefore a defensible reconstruction. Podomere reduction is common and achieved in numerous, lineage-specific ways, even in extant groups (Klementz et al. 2025). It is entirely possible that some of the discrepancies observed in limb patterning of chelicerates, myriapods, and pancrustaceans reflect independent evolutionary paths to arrive at reduced podomere counts, rather than simply developmental system drift alone.

## Conclusion

Testing hypotheses of homology requires both broad taxon sampling and broad sampling of gene expression domains, in pursuit of falsifiability and defensibility in hypothesis-testing. Evolutionary scenarios and homology schemes must integrate and reconcile different lines of evidence, even if these are developed in independent disciplines (*contra* Sharma et al. 2021). Phylogenetically significant lineages, such as sea spiders and centipedes, serve as the linchpins for polarizing ancestral state reconstructions and interpretations of putative derived states, and must therefore receive priority for development of functional genetic tools. Now, given several examples of developmental system drift, such analyses will benefit from the increased emphasis on unbiased approaches to evolutionary developmental biology, rather than the classical candidate gene approaches (Vieira and Nunes-da-Fonseca, 2026). Such efforts may be able to demonstrate how traditional candidate genes are rewired or redeployed within the context of broader gene regulatory networks (Akaiwa et al., 2025; Medina-Jiménez et al., 2026, 2024).

## Acknowledgements

We are indebted to Sarah Swanson and the Newcomb Imaging Center (Department of Biology, University of Wisconsin-Madison) for providing imaging infrastructure and essential training. Alina Farrell assisted with the care and maintenance of *P. opilio*. Confocal microscopy of *P. litorale* was in part performed at the Core Facility Cell Imaging and Ultrastructure Research, University of Vienna – member of the Vienna Life-Science Instruments (VLSI). Discussions with Emily Setton refined an earlier draft of this manuscript. This research was funded in part by the Austrian Science Fund (FWF) 10.55776/PAT1457924 to G.B. and by intramural University of Wisconsin-Madison funds from the Vilas Foundation to P.P.S.

## Author Contributions

**Benjamin Klementz:** conceptualization; methodology; investigation; data curation; writing – original draft; visualization. **Max Hämmerle**: investigation; writing – review and editing; visualization. **Ethan Laumer**: investigation; writing – review and editing. **Sophie Neu**: investigation; writing – review and editing. **Madeline Koebrick:** investigation; writing – review and editing. **Nithya Atla:** investigation; writing – review and editing. **Pranil Panda:** investigation; writing – review and editing. **Neeharika Verma:** resources; writing – review and editing. **Georg Brenneis:** conceptualization; methodology; investigation; data curation; writing – review and editing; visualization; supervision; funding acquisition. **Prashant Sharma:** conceptualization; methodology; investigation; data curation; writing – review and editing; visualization; supervision; funding acquisition.

## Notes

### Competing Interest Statement

The authors have declared no competing interest.

## References

Aase-Remedios, M.E., Janssen, R., Leite, D.J., Sumner-Rooney, L., McGregor, A.P. 2023. Evolution of the spider homeobox gene repertoire by tandem and whole genome duplication. Mol. Biol. Evol. 40, msad239. 10.1093/molbev/msad239.

Abzhanov, A., Kaufman, T.C., 2000. Homologs of Drosophila Appendage Genes in the Patterning of Arthropod Limbs. Dev. Biol. 227, 673–689. 10.1006/dbio.2000.9904

Adamowicz, S.J., Purvis, A., Wills, M.A., 2008. Increasing morphological complexity in multiple parallel lineages of the Crustacea. Proc. Natl. Acad. Sci. 105, 4786–4791. 10.1073/pnas.0709378105

Akaiwa, T., Oda, H., Akiyama-Oda, Y., 2025. Genome-wide quantitative dissection of an arthropod segmented body plan at single-cell resolution. Commun. Biol. 8, 913. 10.1038/s42003-025-08335-x

Akam, M., 1995. Hox Genes and the Evolution of Diverse Body Plans. Phil. Trans.: Biol. Sci. 349, 313–319.

Anderson, L.I., Trewin, N.H., 2003. An Early Devonian arthropod fauna from the Windyfield cherts, Aberdeenshire, Scotland. Palaeontology 46, 467–509. 10.1111/1475-4983.00308

Angelini, D.R., Smith, F.W., Jockusch, E.L., 2012. Extent With Modification: Leg Patterning in the Beetle Tribolium castaneum and the Evolution of Serial Homologs. G3: Genes Genomes Genet. 2, 235–248. 10.1534/g3.111.001537

Arango, C.P., Wheeler, W.C. 2007. Phylogeny of the sea spiders (Arthropoda, Pycnogonida) based on direct optimization of six loci and morphology. Cladistics 23, 255–293. 10.1111/j.1096-0031.2007.00143.x

Bamber, R.N. 2010. Sea-spiders (Pycnogonida) of the north-east Atlantic: keys and notes for the identification of the species, 2nd ed., Synopses of the British Fauna. Field Studies Council, Shrewsbury, UK.

Bicknell, R.D.C., Smith, P.M., Poschmann, M., 2020. Re-evaluating evidence of Australian eurypterids. Gondwana Res. 86, 164–181. 10.1016/j.gr.2020.06.002

Bonatto Paese, C.L., Leite, D.J., Schönauer, A., McGregor, A.P., Russell, S., 2018. Duplication and expression of Sox genes in spiders. BMC Evol. Biol. 18, 205. 10.1186/s12862-018-1337-4

Bond, J.E., Opell, B.D., 1998. Testing Adaptive Radiation and Key Innovation Hypotheses in Spiders. Evolution 52, 403–414. 10.1111/j.1558-5646.1998.tb01641.x

Boxshall, G.A., 2004. The evolution of arthropod limbs. Biol. Rev. 79, 253–300. 10.1017/S1464793103006274

Brenneis, G., Frankowski, K., Maaß, L., Scholtz, G. 2023. The sea spider Pycnogonum litorale overturns the paradigm of the absence of axial regeneration in molting animals. Proc. Natl. Acad. Sci. 120, e2217272120. 10.1703/pnas.2217272120

Brenneis, G., Scholtz, G. 2021. A postlarval instar of Phoxichilidium femoratum (Pycnogonida, Phoxichilidiidae) with an exceptional malformation. J. Morphol. 282, 278–290. 10.1002/jmor.21303

Brenneis, G., Schwentner, M., Giribet, G., Beltz, B.S. 2021. Insights into the genetic regulatory network underlying neurogenesis in the parthenogenetic marbled crayfish Procambarus virginalis. Dev. Neurobiol. 81, 939–974. 10.1002/dneu.22852

Browne, W.E., Price, A.L., Gerberding, M., Patel, N.H., 2005. Stages of embryonic development in the amphipod crustacean, Parhyale hawaiensis. Genesis 42, 124–149. 10.1002/gene.20145

Bruce, H.S., Jerz, G., Kelly, S., McCarthy, J., Pomerantz, A., Senevirathne, G., Sherrard, A., Sun, D.A., Wolff, C., Patel, N.H., 2021. Hybridization Chain Reaction (HCR) In Situ Protocol. protocols.io. 10.17504/protocols.io.bunznvf6

Bruce, H.S., Patel, N.H., 2020. Knockout of crustacean leg patterning genes suggests that insect wings and body walls evolved from ancient leg segments. Nat. Ecol. Evol. 4, 1703–1712. 10.1038/s41559-020-01349-0

Bruce, H.S. 2021. How to align arthropod legs. bioRxiv. 10.1101/2021.01.20.427514

Cano-Sánchez, E., López-González, P.J., 2016. Basal articulation of the palps and ovigers in Antarctic Colossendeis (Pycnogonida; Colossendeidae). Helgol Mar. Res. 70, 22. 10.1186/s10152-016-0474-7

Coulcher, J.F., Edgecombe, G.D., Telford, M.J., 2015. Molecular developmental evidence for a subcoxal origin of pleurites in insects and identity of the subcoxa in the gnathal appendages. Sci. Rep. 5, 15757. 10.1038/srep15757

Crémazy, F., Berta, P., Girard, F., 2001. Genome-wide analysis of Sox genes in Drosophila melanogaster. Mech. Dev. 109, 371–375. 10.1016/S0925-4773(01)00529-9

Crémazy, F., Berta, P., Girard, F., 2000. Sox Neuro, a new Drosophila Sox gene expressed in the developing central nervous system. Mech. Dev. 93, 215–219. 10.1016/S0925-4773(00)00268-9

Dolezal, D.M., Joiner, M.A., Eberl, D.F., 2024. Two distinct functions of Lim1 in the Drosophila antenna. microPub. Biol. 10.17912/micropub.biology.001229

Dong, P.D.S., Chu, J., Panganiban, G., 2001. Proximodistal domain specification and interactions in developing Drosophila appendages. Development 128, 2365–2372. 10.1242/dev.128.12.2365

Dudley, R., Pass, G., 2018. Wings and powered flight: Core novelties in insect evolution. Arthropod Struct. Dev. 47, 319–321. 10.1016/j.asd.2018.06.006

Edgecombe, G.D., Strullu-Derrien, C., Góral, T., Hetherington, A.J., Thompson, C., Koch, M., 2020. Aquatic stem group myriapods close a gap between molecular divergence dates and the terrestrial fossil record. Proc. Natl. Acad. Sci. 117, 8966–8972. 10.1073/pnas.1920733117

Frankowski, K., Miyazaki, K., Brenneis, G. 2022. A microCT-based atlas of the central nervous system and midgut in sea spiders (Pycnogonida) sheds first light on evolutionary trends at the family level. Front. Zool. 19, 14. 10.1186/s12983-022-00459-8

Fu, D., Legg, D.A., Daley, A.C., Budd, G.E., Wu, Y., Zhang, X., 2022. The evolution of biramous appendages revealed by a carapace-bearing Cambrian arthropod. Phil. Trans. R. Soc. B 377, 20210034. 10.1098/rstb.2021.0034

Gainett, G., Crawford, A.R., Klementz, B.C., So, C., Baker, C.M., Setton, E.V.W., Sharma, P.P., 2022. Eggs to long-legs: embryonic staging of the harvestman Phalangium opilio (Opiliones), an emerging model arachnid. Front. Zool. 19, 11. 10.1186/s12983-022-00454-z

Gainett, G., González, V.L., Ballesteros, J.A., Setton, E.V.W., Baker, C.M., Barolo Gargiulo, L., Santibáñez-López, C.E., Coddington, J.A., Sharma, P.P., 2021. The genome of a daddy-long-legs (Opiliones) illuminates the evolution of arachnid appendages. Proc. Roy. Soc. B: Biol. Sci. 288, 20211168. 10.1098/rspb.2021.1168

Gainett, G., Klementz, B.C., Blaszczyk, P.O., Bruce, H.S., Patel, N.H., Sharma, P.P., 2023. Dual Functions of labial Resolve the Hox Logic of Chelicerate Head Segments. Mol. Biol. Evol. 40, msad037. 10.1093/molbev/msad037

García-Bellido, D.C., Collins, D., 2007. Reassessment of the Genus Leanchoilia (Arthropoda, Arachnomorpha) from the Middle Cambrian Burgess Shale, British Columbia, Canada. Palaeontology 50, 693–709. 10.1111/j.1475-4983.2007.00649.x

Gout, J.-F., Hao, Y., Johri, P., Arnaiz, O., Doak, T.G., Bhullar, S., Couloux, A., Guérin, F., Malinsky, S., Potekhin, A., Sawka, N., Sperling, L., Labadie, K., Meyer, E., Duharcourt, S., Lynch, M., 2023. Dynamics of Gene Loss following Ancient Whole-Genome Duplication in the Cryptic Paramecium Complex. Mol. Biol. Evol. 40, msad107. 10.1093/molbev/msad107

Guillermin, C., Tribollet, V., Bouchet, M., Laurencon, A., Zhou, D., Sarnataro, S., Gilquin, L., Stevant, I., Marcy, G., Texeraud, E., Ghavi-Helm, Y., Gillet, B., Hughes, S., Vonau, S., Enriquez, J., 2026. Spatiotemporal control of myoblast identity drives muscle diversity in the Drosophila leg. Sci. Adv. 12, eaed0910. 10.1126/sciadv.aed0910

Hasson, P., DeLaurier, A., Bennett, M., Grigorieva, E., Naiche, L.A., Papaioannou, V.E., Mohun, T.J., Logan, M.P.O., 2010. Tbx4 and Tbx5 Acting in Connective Tissue Are Required for Limb Muscle and Tendon Patterning. Dev. Cell 18, 148–156. 10.1016/j.devcel.2009.11.013

Heingård, M., Turetzek, N., Prpic, N.-M., Janssen, R., 2019. FoxB, a new and highly conserved key factor in arthropod dorsal–ventral (DV) limb patterning. EvoDevo 10, 28. 10.1186/s13227-019-0141-6

Holland, P.W.H., Marlétaz, F., Maeso, I., Dunwell, T.L., Paps, J., 2017. New genes from old: asymmetric divergence of gene duplicates and the evolution of development. Phil. Trans. R. Soc. B 372, 20150480. 10.1098/rstb.2015.0480

Izquierdo-López, A., Caron, J.-B. 2024. The Cambrian Odaraia alata and the colonization of nektonic suspension-feeding niches by early mandibulates. Proc. R. Soc. B 291, 20240622.

Izquierdo-López, A., Caron, J.-B., 2022. Extreme multisegmentation in a giant bivalved arthropod from the Cambrian Burgess Shale. iScience 25, 104675. 10.1016/j.isci.2022.104675

Izquierdo-López, A., Caron, J.-B., 2021. A Burgess Shale mandibulate arthropod with a pygidium: a case of convergent evolution. Pap. Palaeontol. 7, 1877–1894. 10.1002/spp2.1366

Janssen, R., Andersson, E., Betnér, E., Bijl, S., Fowler, W., Höök, L., Leyhr, J., Mannelqvist, A., Panara, V., Smith, K., Tiemann, S., 2018. Embryonic expression patterns and phylogenetic analysis of panarthropod sox genes: insight into nervous system development, segmentation and gonadogenesis. BMC Evol. Biol. 18, 88. 10.1186/s12862-018-1196-z

Klementz, B., Neu, S., Laumer, E., Hinne, I., Setton, E., Verma, N., Hämmerle, M., Rallis, J., Barnett, A., Brenneis, G., Gulia-Nuss, M., Pavlopoulos, A., Sharma, P., 2026. Developmental system drift in the patterning of the arthropod tarsus. Proc. Roy. Soc. B: Biol. Sci. 293, 20252557. 10.1098/rspb.2025.2557

Klementz, B.C., Brenneis, G., Hinne, I.A., Laumer, E.M., Neu, S.M., Hareid, G.M., Gainett, G., Setton, E.V.W., Simian, C., Vrech, D.E., Joyce, I., Barnett, A.A., Patel, N.H., Harvey, M.S., Peretti, A.V., Gulia-Nuss, M., Sharma, P.P., 2024. A Novel Expression Domain of extradenticle Underlies the Evolutionary Developmental Origin of the Chelicerate Patella. Mol. Biol. Evol. 41, msae188. 10.1093/molbev/msae188

Klementz, B.C., Brenneis, G., Laumer, E.M., Neu, S.M., Harvey, M.S., Sharma, P.P., 2025. Evolution and homology of leg segments in Chelicerata: Evo-devo solutions to century-old challenges. Arthropod Struct. Dev. 87, 101446. 10.1016/j.asd.2025.101446

Kojima, T., 2017. Developmental mechanism of the tarsus in insect legs. Curr. Opin. Insect Sci. 19, 36–42. 10.1016/j.cois.2016.11.002

Königsmann, T., Turetzek, N., Pechmann, M., Prpic, N.-M., 2017. Expression and function of the zinc finger transcription factor Sp6–9 in the spider Parasteatoda tepidariorum. Dev. Genes Evol. 227, 389–400. 10.1007/s00427-017-0595-2

Kuehn, E., Clausen, D.S., Null, R.W., Metzger, B.M., Willis, A.D., Özpolat, B.D., 2022. Segment number threshold determines juvenile onset of germline cluster expansion in Platynereis dumerilii. J. Exp. Zool. Part B: Mol. Dev. Evol. 338, 225–240. 10.1002/jez.b.23100

Lamsdell, J.C., 2025. Codex Eurypterida: A Revised Taxonomy Based on Concordant Parsimony and Bayesian Phylogenetic Analyses. Bull. Am. Mus. Nat. Hist. 2025, 1–196. 10.1206/0003-0090.473.1.1

Lamsdell, J.C., Briggs, D.E.G., Liu, H.P., Witzke, B.J., McKay, R.M., 2015. The oldest described eurypterid: a giant Middle Ordovician (Darriwilian) megalograptid from the Winneshiek Lagerstätte of Iowa. BMC Evol. Biol. 15, 169. 10.1186/s12862-015-0443-9

Lewis, E.B., 1978. A gene complex controlling segmentation in Drosophila. Nature 276, 565–570. 10.1038/276565a0

Luo, Y., Chen, N., Gao, Y., Luo, W., Liu, Shanshan, Li, Y., Wang, H., Gao, X., Chen, J., Ye, H., King-Jones, K., Palli, S.R., Li, S., Liu, Suning, 2026. Single-cell transcriptomics of the Drosophila ring gland identifies the SoxN-Vvl complex as a key regulator of juvenile hormone biosynthesis. Proc. Natl. Acad. Sci. 123, e2520504123. 10.1073/pnas.2520504123

Moritz, M. 1957. Embryonalentwicklung der Phalangiiden (Opiliones, Palpatores) unter besonder Berücksichtigung der äußeren Morphologie, der Bildung des Mitteldarmes und der Genitalanlage. Zool. Jahrb. Abt. Anat. Ontog. Tiere 76, 331–370.

Martin, A., Serano, J.M., Jarvis, E., Bruce, H.S., Wang, J., Ray, S., Barker, C.A., O’Connell, L.C., Patel, N.H., 2016. CRISPR/Cas9 Mutagenesis Reveals Versatile Roles of Hox Genes in Crustacean Limb Specification and Evolution. Curr. Biol. 26, 14–26. 10.1016/j.cub.2015.11.021

Medina-Jiménez, B.I., Budd, G.E., Janssen, R., 2026. Single-cell sequencing reveals potential novel insights into appendage-patterning and joint-development in a spider. Dev. Dyn. 255, 671–697. 10.1002/dvdy.70069

Medina-Jiménez, B.I., Budd, G.E., Janssen, R., 2024. Single-cell RNA sequencing of mid-to-late stage spider embryos: new insights into spider development. BMC Genomics 25, 150. 10.1186/s12864-023-09898-x

Ortega-Hernández, J. 2026. A decade after Deuteropoda - insights into the systematics, evolution and dynamics of Cambrian euarthropods. J. Syst. Palaeontol. 24, 2663907. 10.1080/14772019.2026.2663907.

Panganiban, G., Irvine, S.M., Lowe, C., Roehl, H., Corley, L.S., Sherbon, B., Grenier, J.K., Fallon, J.F., Kimble, J., Walker, M., Wray, G.A., Swalla, B.J., Martindale, M.Q., Carroll, S.B., 1997. The origin and evolution of animalnlappendages. Proc. Natl. Acad. Sci. 94, 5162–5166. 10.1073/pnas.94.10.5162

Panganiban, G., Nagy, L., Carroll, S.B., 1994. The role of the Distal-less gene in the development and evolution of insect limbs. Curr. Biol. 4, 671–675. 10.1016/S0960-9822(00)00151-2

Panganiban, G., Sebring, A., Nagy, L., Carroll, S., 1995. The Development of Crustacean Limbs and the Evolution of Arthropods. Science 270, 1363–1366. 10.1126/science.270.5240.1363

Pioro, H.L., Stollewerk, A., 2006. The expression pattern of genes involved in early neurogenesis suggests distinct and conserved functions in the diplopod Glomeris marginata. Dev. Genes Evol. 216, 417–430. 10.1007/s00427-006-0078-3

Prpic, N.-M., Damen, W.G.M., 2004. Expression patterns of leg genes in the mouthparts of the spider Cupiennius salei (Chelicerata: Arachnida). Dev. Genes Evol. 214, 296–302. 10.1007/s00427-004-0393-5

Pueyo, J.I., Galindo, M.I., Bishop, S.A., Couso, J.P., 2000a. Proximal-distal leg development in Drosophila requires the apterous gene and the Lim1 homologue dlim1. Development 127, 5391–5402. 10.1242/dev.127.24.5391

Rizzo, N.P., Bejsovec, A., 2017. SoxNeuro and Shavenbaby act cooperatively to shape denticles in the embryonic epidermis of Drosophila. Development 144, 2248–2258. 10.1242/dev.150169

Sabroux, R., Garwood, R.J., Pisani, D., Donoghue, P.C.J., Edgecombe, G.D., 2024. New insights into the Devonian sea spiders of the Hunsrück Slate (Arthropoda: Pycnogonida). PeerJ 12, e17766. 10.7717/peerj.17766

Schwager, E.E., Sharma, P.P., Clarke, T., Leite, D.J., Wierschin, T., Pechmann, M., Akiyama-Oda, Y., Esposito, L., Bechsgaard, J., Bilde, T., Buffry, A.D., Chao, H., Dinh, H., Doddapaneni, H., Dugan, S., Eibner, C., Extavour, C.G., Funch, P., Garb, J., Gonzalez, L.B., Gonzalez, V.L., Griffiths-Jones, S., Han, Y., Hayashi, C., Hilbrant, M., Hughes, D.S.T., Janssen, R., Lee, S.L., Maeso, I., Murali, S.C., Muzny, D.M., Nunes da Fonseca, R., Paese, C.L.B., Qu, J., Ronshaugen, M., Schomburg, C., Schönauer, A., Stollewerk, A., Torres-Oliva, M., Turetzek, N., Vanthournout, B., Werren, J.H., Wolff, C., Worley, K.C., Bucher, G., Gibbs, R.A., Coddington, J., Oda, H., Stanke, M., Ayoub, N.A., Prpic, N.-M., Flot, J.-F., Posnien, N., Richards, S., McGregor, A.P., 2017. The house spider genome reveals an ancient whole-genome duplication during arachnid evolution. BMC Biol. 15, 62. 10.1186/s12915-017-0399-x

Setton, E.V.W., Ballesteros, J.A., Blaszczyk, P.O., Klementz, B.C., Sharma, P.P., 2024. A taxon-restricted duplicate of Iroquois3 is required for patterning the spider waist. PLOS Biol. 22, e3002771. 10.1371/journal.pbio.3002771

Setton, E.V.W., March, L.E., Nolan, E.D., Jones, T.E., Cho, H., Wheeler, W.C., Extavour, C.G., Sharma, P.P., 2017. Expression and function of spineless orthologs correlate with distal deutocerebral appendage morphology across Arthropoda. Dev. Biol. 430, 224–236. 10.1016/j.ydbio.2017.07.016

Setton, E.V.W., Sharma, P.P., 2018. Cooption of an appendage-patterning gene cassette in the head segmentation of arachnids. Proc. Natl. Acad. Sci. 115, E3491–E3500. 10.1073/pnas.1720193115

Sharma, P.P., Ballesteros, J.A., Santibáñez-López, C.E. 2021. What is an “arachnid”? Consensus, consilience, and confirmation bias in the phylogenetics of Chelicerata. Diversity 13, 568. 10.3390/d13110568

Sharma, P.P., Schwager, E.E., Giribet, G., Jockusch, E.L., Extavour, C.G., 2013. Distal-less and dachshund pattern both plesiomorphic and apomorphic structures in chelicerates: RNA interference in the harvestman Phalangium opilio (Opiliones). Evol. Dev. 15, 228–242. 10.1111/ede.12029

Sharma, P.P., Tarazona, O.A., Lopez, D.H., Schwager, E.E., Cohn, M.J., Wheeler, W.C., Extavour, C.G., 2015. A conserved genetic mechanism specifies deutocerebral appendage identity in insects and arachnids. Proc. R. Soc. B 282, 20150698. 10.1098/rspb.2015.0698

Siveter, D.J., Sabroux, R., Briggs, D.E.G., Siveter, D.J., Sutton, M.D. 2023. Newly discovered morphology of the Silurian sea spider Haliestes and its implications. Pap. Palaeontol. 9, e1528. 10.1002/spp2.1528

Stansbury, M.S., Moczek, A.P., 2013. The Evolvability of Arthropods, in: Minelli, A., Boxshall, G., Fusco, G. (Eds.), Arthropod Biology and Evolution: Molecules, Development, Morphology. Springer, Berlin, Heidelberg, pp. 479–493. 10.1007/978-3-642-36160-9_18

Strausfeld, N.J., Andrew, D.R., Hirth, F., 2026. Cambrian origin of the arachnid brain. Curr. Biol. 36, 259. 10.1016/j.cub.2025.12.019

Strausfeld, N.J., Andrew, D.R., Hou, X., Hirth, F., 2025. Brain anatomy of the Cambrian fossil Jianfengia multisegmentalis informs euarthropod phylogeny. Nat. Commun. 16, 7938. 10.1038/s41467-025-62849-w

Tsuji, T., Sato, A., Hiratani, I., Taira, M., Saigo, K., Kojima, T., 2000. Requirements of Lim1, a Drosophila LIM-homeobox gene, for normal leg and antennal development. Development 127, 4315–4323. 10.1242/dev.127.20.4315

Turchyn, N., Chesebro, J., Hrycaj, S., Couso, J.P., Popadić, A., 2011. Evolution of nubbin function in hemimetabolous and holometabolous insect appendages. Dev. Biol. 357, 83–95. 10.1016/j.ydbio.2011.06.014

Turetzek, N., Khadjeh, S., Schomburg, C., Prpic, N.-M., 2017. Rapid diversification of homothorax expression patterns after gene duplication in spiders. BMC Evol. Biol. 17, 168. 10.1186/s12862-017-1013-0

Vannier, J., Aria, C., Taylor, R.S., Caron, J.-B., 2018. Waptia fieldensis Walcott, a mandibulate arthropod from the middle Cambrian Burgess Shale. R. Soc. Open Sci. 5, 172206. 10.1098/rsos.172206

Vieira, J., Nunes-da-Fonseca, R., 2026. From candidate genes to omics: Unbiased approaches reshaping arthropod Evo-Devo. Genet. Mol. Biol. 49, e20250217. 10.1590/1678-4685-GMB-2025-0217

Yang, J., Ortega-Hernández, J., Legg, D.A., Lan, T., Hou, J., Zhang, X., 2018. Early Cambrian fuxianhuiids from China reveal origin of the gnathobasic protopodite in euarthropods. Nat. Commun. 9, 470. 10.1038/s41467-017-02754-z

Zhai, D., Ortega-Hernández, J., Wolfe, J.M., Hou, X., Cao, C., Liu, Y., 2019. Three-Dimensionally Preserved Appendages in an Early Cambrian Stem-Group Pancrustacean. Curr. Biol. 29, 171–177.e1. 10.1016/j.cub.2018.11.060

Zhang, C., Liu, Y., Ortega-Hernández, J., Wolfe, J., Jin, C., Mai, H., Hou, X., Guo, J., Zhai, D., 2021. Differentiated appendages in Isoxys illuminate origin of arthropodization. 10.21203/rs.3.rs-861892/v1

Zhang, N., 2016. Developmental Studies of Appendage Patterning and Formation in Spiders (doctoralThesis). 10.53846/goediss-5988

Zhang, X., Siveter, D.J., Waloszek, D., Maas, A., 2007. An epipodite-bearing crown-group crustacean from the Lower Cambrian. Nature 449, 595–598. 10.1038/nature06138

