## Supplementary File S1 for "Homoplastic evolution of proximal podomere patterning across Arthropoda"

**Supplementary Figures**

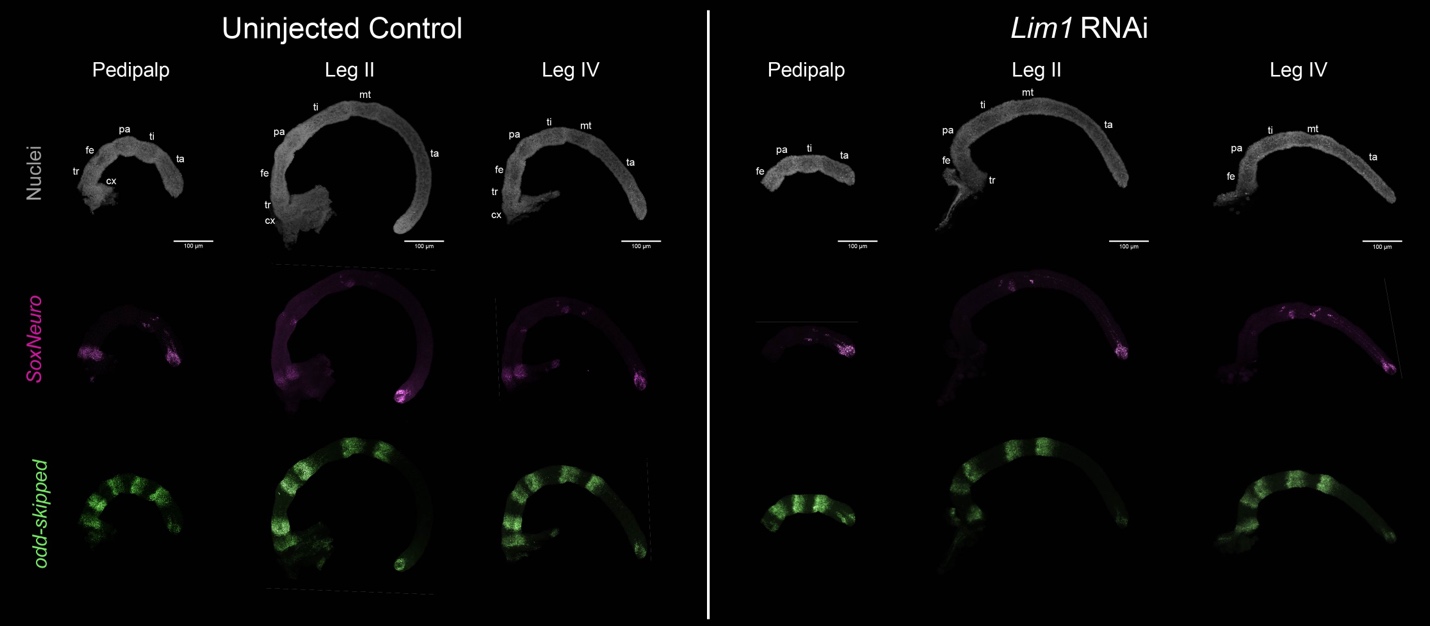

**Fig. S1. Isolated *Po-SoxNeuro* and *Po-odd-skipped* expression following *Po-Lim1* RNAi in *P. opilio*.** All panels represent the same appendages depicted in Fig. 4A-F. Left: appendages dissected from uninjected control embryos. Right: appendages dissected from *Lim1* RNAi embryos. Top row: isolated nuclear counterstaining (grey). Middle row: *Po-SoxNeuro* expression (magenta). Bottom row: *Po-odd-skipped* expression. Abbreviations: cx – coxa; tr – trochanter; fe – femur; pa – patella; ti – tibia; mt – metatarsus; ta – tarsus.

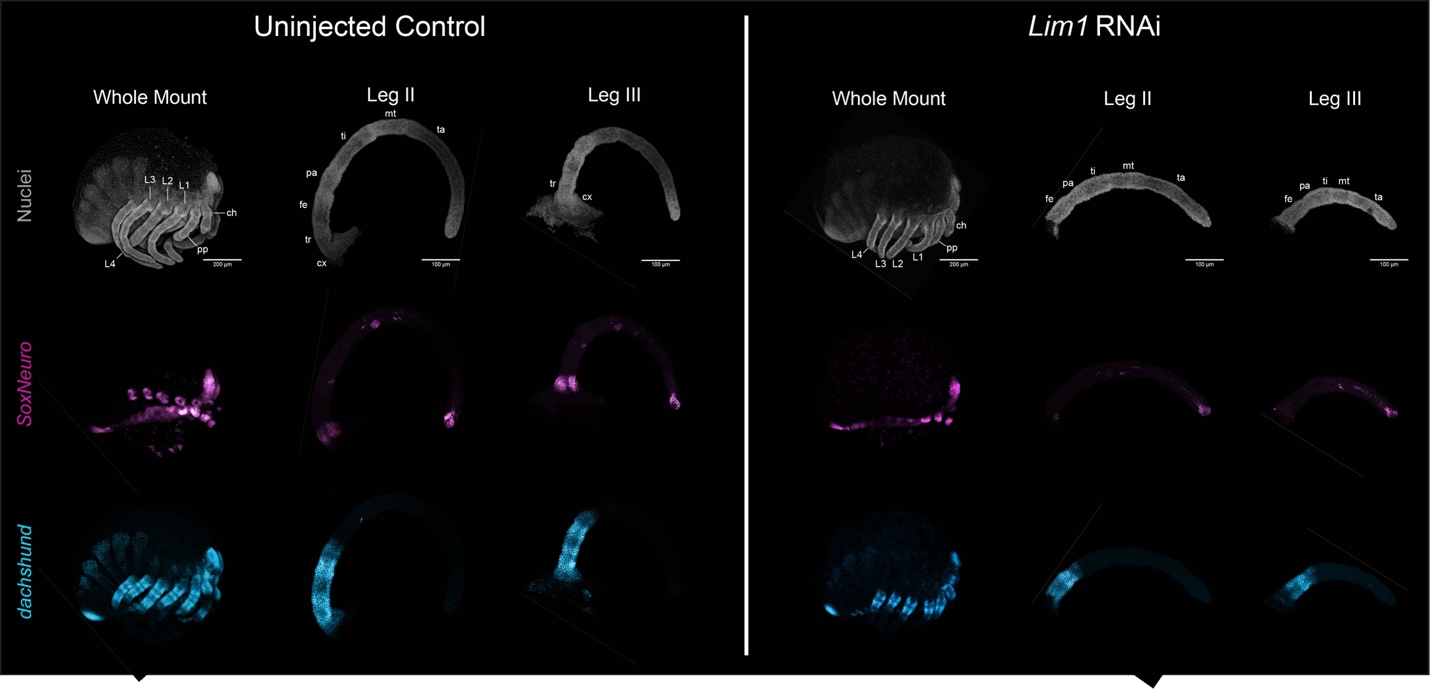

**Fig. S2. Isolated *Po-SoxNeuro* and *Po-dachshund* expression following *Po-Lim1* RNAi in *P. opilio*.** All panels represent the same embryos and appendages depicted in Fig. 4G-L. Left: Uninjected control embryos and appendages. Right: *Lim1* RNAi embryos and appendages. Top row: isolated nuclear counterstaining (grey). Middle row: *Po-SoxNeuro* expression (magenta). Bottom row: *Po-dachshund* expression. Abbreviations: ch – chelicera; pp – pedipalp; L1 – walking leg I; others as in Fig. S1.

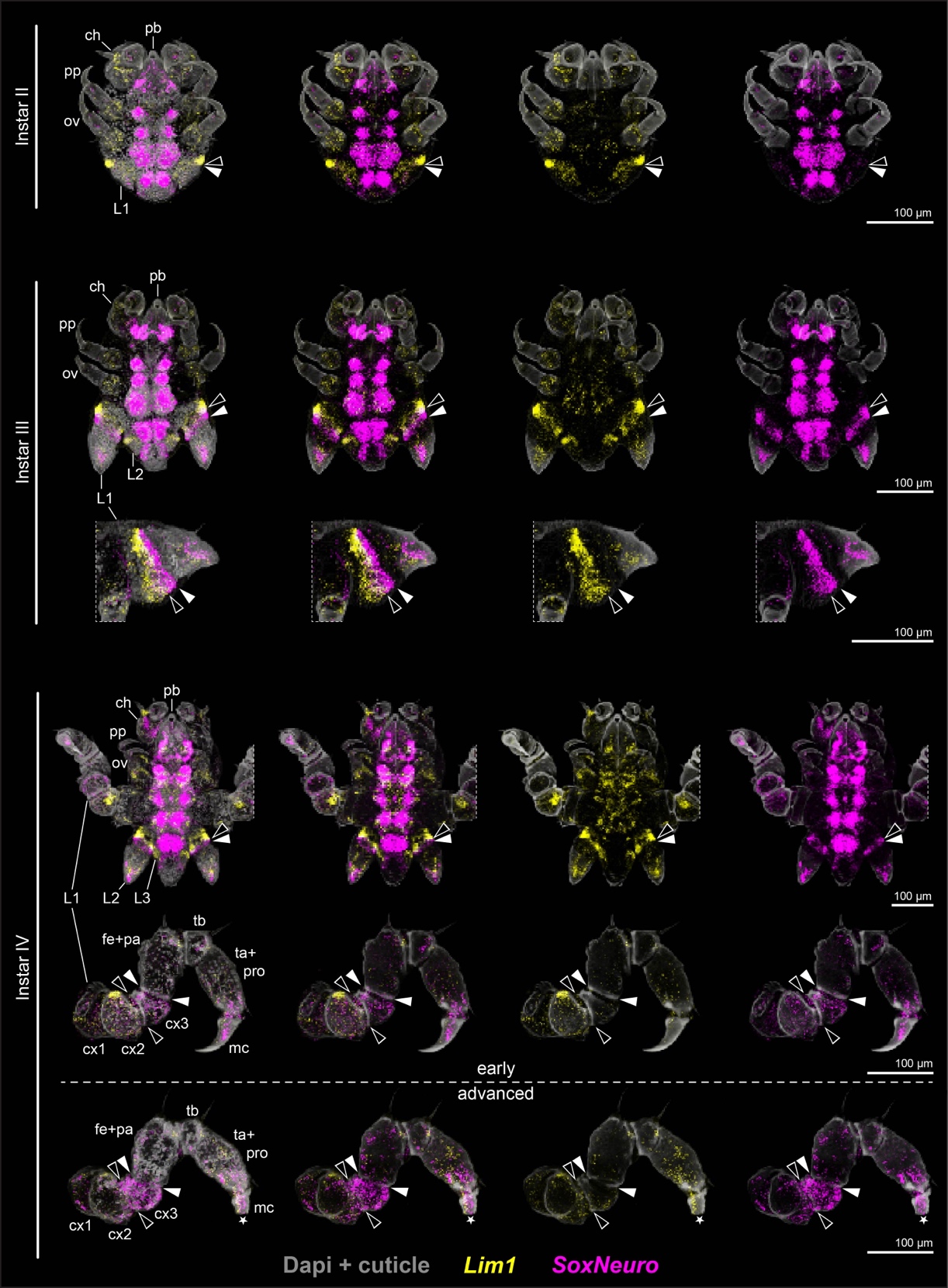

**Fig. S3. Overview of *Pl-Lim1* and *Pl-SoxNeuro* expression during postembryonic development of *P. litorale*.** Whole instars are presented in ventral view and appendages in lateral view. Black arrowhead: distal boundary of appendicular *Pl-Lim1* expression (yellow). White arrowhead: distal boundary of appendicular *Pl-SoxNeuro* expression (magenta). Abbreviations: pb – proboscis; ch – chelifore; ov – oviger; pro – propodus; mc – main claw; others as in Fig. S1.

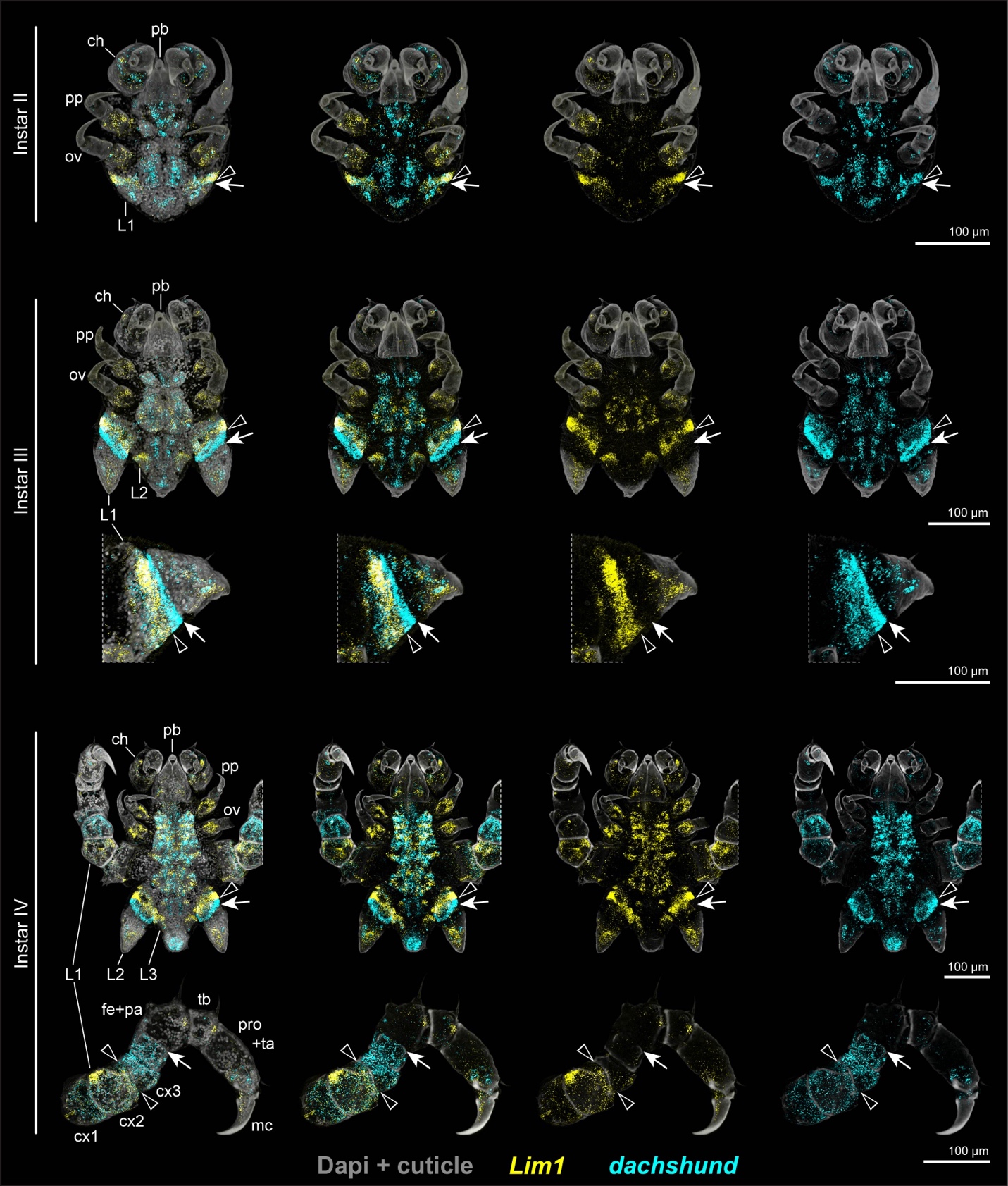

**Fig. S4. Overview of *Pl-Lim1* and *Pl-dachshund* expression during postembryonic development of *P. litorale*.** Whole instars are presented in ventral view and appendages in lateral view. Black arrowhead: distal boundary of appendicular *Pl-Lim1* expression (yellow). White arrow: distal boundary of appendicular *Pl-dachshund* expression (cyan). Abbreviations as in Figs. S1, S3.

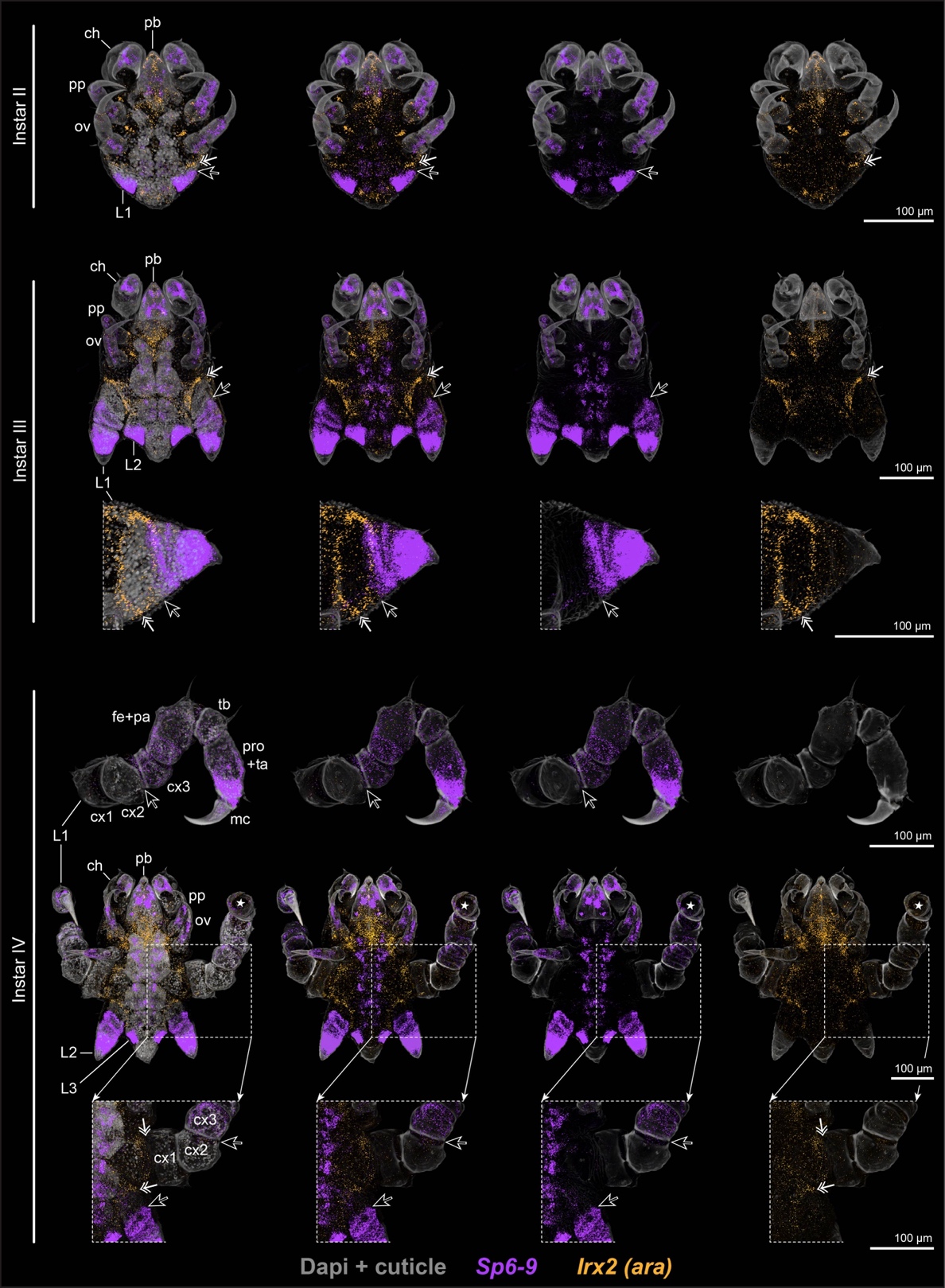

**Fig. S5. Overview of *Pl-Irx2* and *Pl-Sp6-9* expression during postembryonic development of *P. litorale*.** Whole instars are presented in ventral view and appendages in lateral view. Dual white arrow: distal boundary of *Pl-Irx2* expression (orange) in the non-appendicular body wall. Black arrow: proximal boundary of appendicular *Pl-Sp6-9* expression (violet). Damage to the distal territory of one instar IV appendage is indicated by the star. Abbreviations as in Figs. S1, S3.

**Supplementary Materials and Methods**

*Gene Expression Assays*

Probes for *Pycnogonum litorale SoxN* HCR assays were designed by Molecular Instruments using the following CDS template:

>*Plit_SoxNeuro*

TAACATGTGTGTGTGATCTATCTTTGAAAAAATTGGATTCTTGTTTAAAAAAAAGAAAAGATTATTTTTGGATCTCGTGATTGGAATTTAAAGTTTCTTATCTATATGGACCTGTACTCAACCGTCACGCTGTATTCCCCACCGCATTACGTTTCTCTACCTCCATCTGGTATGGAGACGGACTCTTTGAAGGCCGCTGCCCAAATAACACCGGCAGCGTACCCTCATCATCATCATCCGCATCAGACCCATCATGCGTCTGCAGCGCCGCCATCAAATAACACGACACTGCCGCCATCGTCGATTCACGGTGGTCATCATCAAACTCATATTCACAATAGTCCGAATTCCGGTGGTGGAGGAGGTGGAGGTCATCATCATGGTCAACCTGGCAATCCGAATAGCATCCAGAATAAAAACAACCAGAATTCCGATCGTGTTAAGCGGCCCATGAACGCATTTATGGTATGGTCAAGAGGTCAGCGGCGAAAGATGGCTCAAGAGAATCCCAAGATGCATAATTCGGAAATAAGCAAGCGATTGGGCGCGACGTGGAAGTTACTGTCCGAAGCTGAAAAGAGGCCATTCATCGACGAAGCCAAGAGGTTAAGGGCCGTTCACATGAAAGAACATCCCGATTATAAATACAGGCCTAGAAGGAAGACGAAGACGTTAATGAAGAAGGATAAATATCCGTTGACGGCAGCAATGTTACCGAGTGATGGCCGGTCGTCGGGGGTTCAACAGGTCTCTCGCGATGTTTATCATCAGATGAATGGTTATATGCCCAACGGTTATTCGATGATGCATGATGCACATGCTGCTGCCGCTTACCAACACGCTGCCGTAGCGGCCGGCGGTTTCGGTTCTCAAGGAAACGGCGGACTTTATTCGCGTTACGACATGACCCCGTCGCAAATGTCGTCTAGTAGTCCGGCAATGTCATCTTATTTGAACAGTAGTGCATATAATTCGCCGATGGCCGGTATGACAACTTATTCACCCATGAATTCGATGTCTAACGGACAGATTATTAAATCGGAAAGTGGTTCGCCTGTCAGCCCGCCAACATCCGCGCCTCAGATGATGAGGAGACCGGGCTGTCCGAGTGATTTACGTACAGAGATGATTAGCATGTATCTACCTGATGGAACTGCTGATCCTAATGCACAAGCCAGATTAGCTAGTATGGGCCATTACGGCCAATCAGAACATTCTGTTCACGTGCCGCCGTTAACACACATGTAGAACGATATTTTTTAA

**Table S1.** Accession numbers and source publications of genomic or transcriptomic assemblies queried for gene homologs assayed in HCR expression assays.

| **Taxon** | **NCBI Accession Number** | **Assembly Type** | **Source Publication** |
| --- | --- | --- | --- |
| *Phalangium opilio* | GCA_054372755.1 | Genome | Aase-Remedios et al. (2025) |
| *Pycnogonum litorale* | GCA_964442445.1 | Genome | Papadopoulos et al. (2025) |
| *Lithobius atkinsoni* | SAMN10093766 | Transcriptome | Setton and Sharma (2018) |
| *Parhyale hawaiensis* | GCA_001587735.2 | Genome | Kao et al. (2016) |

**Table S2.** Probe pairs designed for *Phalangium opilio Sp6-9* HCR assays (B1 initiator).

| **Pair** | **Initiator** | **Spacer** | **Hybridization** | **Hybridization** | **Spacer** | **Initiator** |
| --- | --- | --- | --- | --- | --- | --- |
| 1 | GAGGAGGGCAGCAAACGG | AA | CATAGCGGTGTAATAGTGGTGTCGC | TCACGACTGTCGAATGTCATGAAGA | TA | GAAGAGTCTTCCTTTACG |
| 2 | GAGGAGGGCAGCAAACGG | AA | CTTGAATTCTCAGCCTGAGGTCGGC | TGAAGACTCGGGAAGTCGTGATTGT | TA | GAAGAGTCTTCCTTTACG |
| 3 | GAGGAGGGCAGCAAACGG | AA | GTCCATCAGCCAGGCCCAAAGGTCC | TTATTTTCGGTTTGCGATCAACTGG | TA | GAAGAGTCTTCCTTTACG |
| 4 | GAGGAGGGCAGCAAACGG | AA | GCGTTCCGACGTTATTGGGTGAATT | GGGGTCCGCCACCGGAGATTAAGGG | TA | GAAGAGTCTTCCTTTACG |
| 5 | GAGGAGGGCAGCAAACGG | AA | TTCGGAGTCGCTGCAAGTTCCACTG | AGTATGAGGTTCGCTCTGACTGTCG | TA | GAAGAGTCTTCCTTTACG |
| 6 | GAGGAGGGCAGCAAACGG | AA | CTTCCGCTGTGCGTCTTAACGTGTT | CTGCCTTTCTTGCCGCCTTCTCCGC | TA | GAAGAGTCTTCCTTTACG |
| 7 | GAGGAGGGCAGCAAACGG | AA | TATTGCAAACGGGGCAAGCGAAACG | ACAAATGGTCGGACCTCATGAAACG | TA | GAAGAGTCTTCCTTTACG |
| 8 | GAGGAGGGCAGCAAACGG | AA | TTGCAATTCGTCCGAACGCGTGAAA | CTCTCCGGTATGCGTCCTCAAATGA | TA | GAAGAGTCTTCCTTTACG |
| 9 | GAGGAGGGCAGCAAACGG | AA | CTCTCCGGTGTGCCATCGTAAATGG | AAAAAGCCAATTGCAGACGAAAGGC | TA | GAAGAGTCTTCCTTTACG |
| 10 | GAGGAGGGCAGCAAACGG | AA | TTGCCGCACCCGGGTATGTGGCAAC | TTCAAATGCGAAGTTTTGCCGTAAA | TA | GAAGAGTCTTCCTTTACG |
| 11 | GAGGAGGGCAGCAAACGG | AA | CGCAAGTGGCCCTTCCGGTGTATCT | GCTCCGCTTCTTGGCAGTTGGGGCA | TA | GAAGAGTCTTCCTTTACG |
| 12 | GAGGAGGGCAGCAAACGG | AA | TGTCCCGTCGCGCTCAGAGAAGACT | TGATGGGCGTGATGGTGAGCCGGAT | TA | GAAGAGTCTTCCTTTACG |
| 13 | GAGGAGGGCAGCAAACGG | AA | AGACTTGAGACGTTGGGAGCATGGA | GCGACAAGCCGAACGGTCCCTGAGA | TA | GAAGAGTCTTCCTTTACG |
| 14 | GAGGAGGGCAGCAAACGG | AA | GGAGGATAATAAATGCGCGGCTGCC | GTACGTGTCTTGTAGCAAATGTTGG | TA | GAAGAGTCTTCCTTTACG |
| 15 | GAGGAGGGCAGCAAACGG | AA | GGCCCGTAATTGACTTGCGAATGAA | GGACCCAGGCCGTAAGCCTCGGCCG | TA | GAAGAGTCTTCCTTTACG |
| 16 | GAGGAGGGCAGCAAACGG | AA | TGTTGAGCGTATTGACCGAGGAAGA | CCGCGTGGACATCCCACCAGGAAGC | TA | GAAGAGTCTTCCTTTACG |
| 17 | GAGGAGGGCAGCAAACGG | AA | ACTCTCGTAAGGATGCGGCGCGTGA | TTCCGGTTTAATGGCGTTGAAGGCC | TA | GAAGAGTCTTCCTTTACG |
| 18 | GAGGAGGGCAGCAAACGG | AA | GACTGATTCGAAGCCGCGTCGGAGA | CGCGAATAGACCGAGCCCAAATGCG | TA | GAAGAGTCTTCCTTTACG |
| 19 | GAGGAGGGCAGCAAACGG | AA | CGTGTTGCTCAGCGAACTCGAAGAA | GCGGTTGTAACCCGCCAGCGAAACG | TA | GAAGAGTCTTCCTTTACG |
| 20 | GAGGAGGGCAGCAAACGG | AA | GAATGCGAAGAAGGTTGATGATTAT | TTAGCGGAAGAAACGCGTTGAAGTG | TA | GAAGAGTCTTCCTTTACG |
| 21 | GAGGAGGGCAGCAAACGG | AA | CCCTGAATGCTCCCCGAACTGTTGT | GCTCCGGGACTCGAGGCTTGTTGTC | TA | GAAGAGTCTTCCTTTACG |
| 22 | GAGGAGGGCAGCAAACGG | AA | GGCTAACGGAGGCGGACTTTTGCTA | GTGGAATCCTTTGCCGACTGCGGCG | TA | GAAGAGTCTTCCTTTACG |

**Table S3.** Probe pairs designed for *Parhyale hawaiensis Lim-1* HCR assays (B2 initiator).

| **Pair** | **Initiator** | **Spacer** | **Hybridization** | **Hybridization** | **Spacer** | **Initiator** |
| --- | --- | --- | --- | --- | --- | --- |
| 1 | CCTCGTAAATCCTCATCA | AA | CCGGTCATTGAAATTCTGGGTACTG | TTACCACGCAAGGATCTCACTGTTC | AA | ATCATCCAGTAAACCGCC |
| 2 | CCTCGTAAATCCTCATCA | AA | ATGGGTGGAGGCTGCTGAACAAATG | GGGAGCGTGACGTCATCTGGAGATG | AA | ATCATCCAGTAAACCGCC |
| 3 | CCTCGTAAATCCTCATCA | AA | TCATGGGCCCACCGGCCGGTAGTGG | GGGGCATTCCAGGGAAGTCTGTGGC | AA | ATCATCCAGTAAACCGCC |
| 4 | CCTCGTAAATCCTCATCA | AA | ACCCATTGGAGGCGGCTGCTGATGG | CTCTAAGACTTGGCCCATATTAGGG | AA | ATCATCCAGTAAACCGCC |
| 5 | CCTCGTAAATCCTCATCA | AA | GATCATCCATGCCGAGAGGGAGGTC | CCCCGTAGCCCGGGAAGAAGCCATA | AA | ATCATCCAGTAAACCGCC |
| 6 | CCTCGTAAATCCTCATCA | AA | CATCTGGGATAGTTGCTTCATGCGA | GGGGTTGAAGAGGTGTGGTCTCATC | AA | ATCATCCAGTAAACCGCC |
| 7 | CCTCGTAAATCCTCATCA | AA | ACCTGGATGACCCGCATGGGCAGCC | TCTTTGCTGCGCTTGTTTTGGAACC | AA | ATCATCCAGTAAACCGCC |
| 8 | CCTCGTAAATCCTCATCA | AA | GTCGGGTCGGCTTCGGTGTGTTGTT | TTTCCTTAGCTAGCTGCTCTCTGAT | AA | ATCATCCAGTAAACCGCC |
| 9 | CCTCGTAAATCCTCATCA | AA | AGCTTTGATGGTGGTGCGGGGCCCC | TGCTACTTTGAGCACCTCGAGCTGC | AA | ATCATCCAGTAAACCGCC |
| 10 | CCTCGTAAATCCTCATCA | AA | GGAACTTTGGCTCCAGCAGTGGTGG | CTCTTGTTGCCTCCCTTCTCATCAT | AA | ATCATCCAGTAAACCGCC |
| 11 | CCTCGTAAATCCTCATCA | AA | CGGAGACGTCGGAGTCGTGAAATGG | CTTTGGACTCCGAGCTCGCCTCATT | AA | ATCATCCAGTAAACCGCC |
| 12 | CCTCGTAAATCCTCATCA | AA | GCTGTTTACACTCAGACTGTAAGAT | TCCAGGTTCCTTTTCCGGGGTCGCT | AA | ATCATCCAGTAAACCGCC |
| 13 | CCTCGTAAATCCTCATCA | AA | GGAGCCAGGGAGTCGTGTAAAGAGG | AGGGAAATGTTGGGCTTGTTGTCAG | AA | ATCATCCAGTAAACCGCC |
| 14 | CCTCGTAAATCCTCATCA | AA | TCAGAGGTGTGCTGGGTCTGGCCAC | GTTTGGAGGCAGGAGGAAGCTGTGC | AA | ATCATCCAGTAAACCGCC |
| 15 | CCTCGTAAATCCTCATCA | AA | TGATGCGAGTGATGGTGATGGTAAC | TAGACCTCCATGCAAGAGGGACGGC | AA | ATCATCCAGTAAACCGCC |
| 16 | CCTCGTAAATCCTCATCA | AA | TGAGGTGAATCCAAGTGGGACGTTG | GCATGTAAAGGTGGCAGACTACTGG | AA | ATCATCCAGTAAACCGCC |
| 17 | CCTCGTAAATCCTCATCA | AA | TGTTGTTGGCTTCCAGTAAAGGAGG | AATTATTGTTACTGTGATCGGGATT | AA | ATCATCCAGTAAACCGCC |
| 18 | CCTCGTAAATCCTCATCA | AA | CGTGGCTCCACAGTCTCCTTCAACC | TAGGTCCAGTTCCTCATCTTCTTCG | AA | ATCATCCAGTAAACCGCC |

**Table S4.** Probe pairs designed for *Parhyale hawaiensis SoxNeuro* HCR assays (B3 initiator).

| **Pair** | **Initiator** | **Spacer** | **Hybridization** | **Hybridization** | **Spacer** | **Initiator** |
| --- | --- | --- | --- | --- | --- | --- |
| 1 | GTCCCTGCCTCTATATCT | TT | GGCGTTCATCTGGTCATGCGGATTG | TTACATATGTGTTAACGGCATAGGC | TT | CCACTCAACTTTAACCCG |
| 2 | GTCCCTGCCTCTATATCT | TT | ACGTCATCGAGGCCATCTGAGACAT | AGGCTGAGGATCCCAGAGATCCTAC | TT | CCACTCAACTTTAACCCG |
| 3 | GTCCCTGCCTCTATATCT | TT | AGAACTGCTGTAAGTGCCATTCATG | ATAAGGAGAAGGAGTAGAGGCATAC | TT | CCACTCAACTTTAACCCG |
| 4 | GTCCCTGCCTCTATATCT | TT | TATCCTCGCGTCATGTCGTATCCTC | GATGTTGGACTAGCTGCAGTAGGTG | TT | CCACTCAACTTTAACCCG |
| 5 | GTCCCTGCCTCTATATCT | TT | TTTGACTACCATACGCTGCTGCACT | ACACTTGACCAGTCATATAAGGATA | TT | CCACTCAACTTTAACCCG |
| 6 | GTCCCTGCCTCTATATCT | TT | GTCACCGTTGGGCATATAGCCGTTA | GTATCCATAACCTGCATAGTCGGAG | TT | CCACTCAACTTTAACCCG |
| 7 | GTCCCTGCCTCTATATCT | TT | TTTGGTTTTCCTCCGAGGTTTGTAG | GGGGTACTTGTCCTTCTTCATCAGA | TT | CCACTCAACTTTAACCCG |
| 8 | GTCCCTGCCTCTATATCT | TT | AGTCTCTTCGCTTCATCGATGAACG | GGGTGTTCCTTCATATGTACTGCCC | TT | CCACTCAACTTTAACCCG |
| 9 | GTCCCTGCCTCTATATCT | TT | ACTCCGCTCCCAGCCGTTTAGATAT | TCTTCTCGGATTCCGTTAATAACTT | TT | CCACTCAACTTTAACCCG |
| 10 | GTCCCTGCCTCTATATCT | TT | GGCCATTTTACGTCTCTGCCCTCTG | AGAATTATGCATCTTGGGGTTTTCT | TT | CCACTCAACTTTAACCCG |
| 11 | GTCCCTGCCTCTATATCT | TT | TTGACCCTCTCAGCCGCTGCATTCG | CACACCATAAATGCATTCATCGGCC | TT | CCACTCAACTTTAACCCG |
| 12 | GTCCCTGCCTCTATATCT | TT | ACGTGTTGTTTGGCGTACCATTGTT | CGGCGCTCTTGCTATTCGAGTTGTT | TT | CCACTCAACTTTAACCCG |
| 13 | GTCCCTGCCTCTATATCT | TT | ATTGGAGCTTGGTACGTGTGTCTGC | GTTAGTACTATTATTGTTGTTGTTA | TT | CCACTCAACTTTAACCCG |
| 14 | GTCCCTGCCTCTATATCT | TT | CTACCGAGAGATGGCGGCGGAGAAT | GGAGAATGCGAACTCATGCTTCCTA | TT | CCACTCAACTTTAACCCG |
| 15 | GTCCCTGCCTCTATATCT | TT | AGTGGTTGTGTGCATGGGTGTGATT | TCGGGTGGGTGACATGATGGGACAA | TT | CCACTCAACTTTAACCCG |
| 16 | GTCCCTGCCTCTATATCT | TT | AAGGGAATGTGATGAGTGTGGAGGT | CGTTATGGATGTGTGATGATTATGT | TT | CCACTCAACTTTAACCCG |
| 17 | GTCCCTGCCTCTATATCT | TT | AGGTCTGTGGGGTCCAGCGTGAGCA | TGTGGAGAATGTAAGGCGCTGGTTT | TT | CCACTCAACTTTAACCCG |

**Table S5.** Probe pairs designed for *Lithobius atkinsoni SoxNeuro* HCR assays (B3 initiator).

| **Pair** | **Initiator** | **Spacer** | **Hybridization** | **Hybridization** | **Spacer** | **Initiator** |
| --- | --- | --- | --- | --- | --- | --- |
| 1 | GTCCCTGCCTCTATATCT | TT | ACGAATTGGACGACCCGTTCATGTA | AGTACGGCGACGCCATGGACATGCT | TT | CCACTCAACTTTAACCCG |
| 2 | GTCCCTGCCTCTATATCT | TT | CATTGCCGGCCATCTGGCTACTGAA | CCATGTCGTAGCGAGAGTATAAGGA | TT | CCACTCAACTTTAACCCG |
| 3 | GTCCCTGCCTCTATATCT | TT | TGGGCATGTAGCCGTTCATTTGGTA | GATCGTGCATCATGCTGGGGTAGCC | TT | CCACTCAACTTTAACCCG |
| 4 | GTCCCTGCCTCTATATCT | TT | TCCTTCTTCATGAGGGTCTTAGTCT | AGCAGACCCGTGGCTAGCGGGTACT | TT | CCACTCAACTTTAACCCG |
| 5 | GTCCCTGCCTCTATATCT | TT | TCGGCACCGAGACGCTTGCTGATTT | TTCTCCGATTCGGACAACAGCTTCC | TT | CCACTCAACTTTAACCCG |
| 6 | GTCCCTGCCTCTATATCT | TT | CCATTTTTCGACGCTGGCCTCTTGA | AGTTGTGCATTTTCGGGTTCTCTTG | TT | CCACTCAACTTTAACCCG |
| 7 | GTCCCTGCCTCTATATCT | TT | CACGCGATCGGCGGCGTTATTCTTG | AACCATGAAGGCATTCATGGGACGC | TT | CCACTCAACTTTAACCCG |
| 8 | GTCCCTGCCTCTATATCT | TT | GGCTCACGCCAAAACTTTTAAACGA | GGATCCACACGGCGAAAGCTGAGGG | TT | CCACTCAACTTTAACCCG |
| 9 | GTCCCTGCCTCTATATCT | TT | ACAACGCACGTTGAACAGAAACAAC | GGGGTAGCGGGCAAACCGACCAGCA | TT | CCACTCAACTTTAACCCG |
| 10 | GTCCCTGCCTCTATATCT | TT | TTTTGGGGTGAGTTCCCAATATGGC | TTCGGCTTTTTGAAAACTCACCTTC | TT | CCACTCAACTTTAACCCG |
| 11 | GTCCCTGCCTCTATATCT | TT | ACAGCCCCACCACTGCCACAAAGTT | GTCGCTTGAACAGCCGAGTGTGTTA | TT | CCACTCAACTTTAACCCG |
| 12 | GTCCCTGCCTCTATATCT | TT | CTGAAAAACTCACCACAGTCTCTCA | AAACACAACAAGTCACTTTTGCCAA | TT | CCACTCAACTTTAACCCG |
| 13 | GTCCCTGCCTCTATATCT | TT | GAAGGAGCAGGCAACACACGACTAA | GTCCTTGTGAGCGCAGTTCGGCAGT | TT | CCACTCAACTTTAACCCG |
| 14 | GTCCCTGCCTCTATATCT | TT | GCGCCCTACGCTACGAGCTTATTGA | ACGCTCGCCTTGTTCTTTGCCTACT | TT | CCACTCAACTTTAACCCG |

**Table S6.** Probe pairs designed for *Pycnogonum litorale Lim1* HCR *in situ* hybridization (B3 initiator).

| Pair | Initiator | Spacer | Hybridization | Hybridization | Spacer | Initiator |
| --- | --- | --- | --- | --- | --- | --- |

| **1** | GTCCCTGCCTCTATATCT | tt | CGCACCATCTTGATTCACGTGTAAA | TCGCGAGCCGTTCATACGGTCTAAG | tt | CCACTCAACTTTAACCCG |
| --- | --- | --- | --- | --- | --- | --- |
| **2** | GTCCCTGCCTCTATATCT | tt | CTTAAAGTTCAGAAACGATGAGTTG | AACATCAGTCCAAGACTTCCAAGGA | tt | CCACTCAACTTTAACCCG |
| **3** | GTCCCTGCCTCTATATCT | tt | TTCATTACCGACGGATGAGTGAGCG | ATGTGCCGTCACCATACGGGAGTCT | tt | CCACTCAACTTTAACCCG |
| **4** | GTCCCTGCCTCTATATCT | tt | ATGAGGTTCTCGGAGGAGGTCCGTT | TCGAGGTGAAATTTTCTCCGGATGA | tt | CCACTCAACTTTAACCCG |
| **5** | GTCCCTGCCTCTATATCT | tt | AAGACTGAATCTGGACTAGAATTTC | CCTCCTCCTCCTCCTTCAGGTAAAT | tt | CCACTCAACTTTAACCCG |
| **6** | GTCCCTGCCTCTATATCT | tt | AGTGACCGTCGTTATGATTTACGCC | GTCCGACCATCAGGTCATGCGACAT | tt | CCACTCAACTTTAACCCG |
| **7** | GTCCCTGCCTCTATATCT | tt | TCTGCGGCATGTGTCCTTGCGGAAA | CGGGTGGCACTGGCTGTTCTATACT | tt | CCACTCAACTTTAACCCG |
| **8** | GTCCCTGCCTCTATATCT | tt | ATGTTGTCCAGGATGACCCATACCG | AGGCATTCCATCCGGTAAACCACCG | tt | CCACTCAACTTTAACCCG |
| **9** | GTCCCTGCCTCTATATCT | tt | GGTTGATTACCGTAAACGTCGCCTT | TGTTGACCAGGAAAGAAATCGTAAA | tt | CCACTCAACTTTAACCCG |
| **10** | GTCCCTGCCTCTATATCT | tt | GTGTTGACCCGTGATGAGGTAAATC | TAGCGCCATCTGCAAAGTAACCGAA | tt | CCACTCAACTTTAACCCG |
| **11** | GTCCCTGCCTCTATATCT | tt | AGACATGCCAGGCCTTAAGGCTCTC | ATCGTCTAAACCGCAACCACTATCA | tt | CCACTCAACTTTAACCCG |
| **12** | GTCCCTGCCTCTATATCT | tt | GAGTACTGTTCTCCCTGTTTTCGTG | TTTTACCGCCGACGGCGTTATCATC | tt | CCACTCAACTTTAACCCG |
| **13** | GTCCCTGCCTCTATATCT | tt | AGAATCCTTATCGACCAGAGTCGAG | GGAATGATTGCTATCACCGTCTATA | tt | CCACTCAACTTTAACCCG |
| **14** | GTCCCTGCCTCTATATCT | tt | GTTCACTCTCGTCGCCGTCCGATAA | TCGGAACTGGATTCAGCACACTTCC | tt | CCACTCAACTTTAACCCG |
| **15** | GTCCCTGCCTCTATATCT | tt | AACGGAATTTCTGGTGCTCATGTAA | TGCTTGCTCCTCCGGAGAATTTAGG | tt | CCACTCAACTTTAACCCG |
| **16** | GTCCCTGCCTCTATATCT | tt | CTCCGGTCGACAGTTGTTTCCTACA | TATTATCGTCCAAGACGTATAGTTC | tt | CCACTCAACTTTAACCCG |
| **17** | GTCCCTGCCTCTATATCT | tt | AACTTTATTCTTAGCCCGACGGACG | CATGCACGTGAAGCATTTCAAATGG | tt | CCACTCAACTTTAACCCG |
| **18** | GTCCCTGCCTCTATATCT | tt | GGCACACTTGGTACCGTAACGTCTG | ACTGGGCGAGATGCCTTGCGAACAA | tt | CCACTCAACTTTAACCCG |
| **19** | GTCCCTGCCTCTATATCT | tt | ACACATCGGACGCATTTAGCGTGCC | TTTTCTGTCAGGTGACATTTGCAAT | tt | CCACTCAACTTTAACCCG |
| **20** | GTCCCTGCCTCTATATCT | tt | GGTCTAATATCGGCCTGTCGCAACC | CTCGATCGAGAACGTTGAAAAGAAA | tt | CCACTCAACTTTAACCCG |
| **21** | GTCCCTGCCTCTATATCT | tt | TTGCATAGATATTCTCATTGTCGTC | GCGAATGACTTGGAATGTTATCACT | tt | CCACTCAACTTTAACCCG |

**Table S7.** Probe pairs designed for *Pycnogonum litorale Irx-2* HCR *in situ* hybridization (B1 initiator).

| Pair | Initiator | Spacer | Hybridization | Hybridization | Spacer | Initiator |
| --- | --- | --- | --- | --- | --- | --- |

| **1** | GAGGAGGGCAGCAAACGG | aa | GCTCACCTGATTTCCCGTGGGATTG | ATATATATCAAAGGAGCGAATCTGC | ta | GAAGAGTCTTCCTTTACG |
| --- | --- | --- | --- | --- | --- | --- |
| **2** | GAGGAGGGCAGCAAACGG | aa | AGATTTTCGTTTGCGTCGATAACAC | TAAACAATTTCAACTGCCAGTTCAC | ta | GAAGAGTCTTCCTTTACG |
| **3** | GAGGAGGGCAGCAAACGG | aa | ACGAGTACAGGGTGGCGTTTGCGTT | GATGGAATCGGCGAACTTATTCGAA | ta | GAAGAGTCTTCCTTTACG |
| **4** | GAGGAGGGCAGCAAACGG | aa | CCGTTAAGTCTCGAGCCGAGAACGA | GGAATACCGAAGACGGGAACGTCGA | ta | GAAGAGTCTTCCTTTACG |
| **5** | GAGGAGGGCAGCAAACGG | aa | CCGCTGACCGATTCCAAAACAATAA | ACTAGAAGATCGCGACGCCGACAAC | ta | GAAGAGTCTTCCTTTACG |
| **6** | GAGGAGGGCAGCAAACGG | aa | TTTATCGTCTTTATTGGCGGCCGTG | ACAGAGTCGATTGTTTTCGATTATC | ta | GAAGAGTCTTCCTTTACG |
| **7** | GAGGAGGGCAGCAAACGG | aa | TCTTCGTTACGCGACGACGAAGCGA | GACCAAATCTTCGGTTTGGTCTTGG | ta | GAAGAGTCTTCCTTTACG |
| **8** | GAGGAGGGCAGCAAACGG | aa | AGGAACGATTCCGGCAGTTATTTGG | TCTTCGAAGACGAAATATTTGGCGA | ta | GAAGAGTCTTCCTTTACG |
| **9** | GAGGAGGGCAGCAAACGG | aa | CTGACTAGAATCGTGAATCCGATTT | CGGACAAGACGGCCGATTAATTGGT | ta | GAAGAGTCTTCCTTTACG |
| **10** | GAGGAGGGCAGCAAACGG | aa | CTTGAAGCTGGACCGGCATGATCGA | GCAATAATCTTCCCGAAATCGTCCG | ta | GAAGAGTCTTCCTTTACG |
| **11** | GAGGAGGGCAGCAAACGG | aa | AAAGTCGAAACAGTTGTACCCGTAA | TTTTCGAACCAGAGAAGCACCTCCG | ta | GAAGAGTCTTCCTTTACG |
| **12** | GAGGAGGGCAGCAAACGG | aa | ACTCGTTGGATCAGTCAATCTATCC | TGGCACCGTCATCGGATGATGCCAT | ta | GAAGAGTCTTCCTTTACG |
| **13** | GAGGAGGGCAGCAAACGG | aa | CGAATCGTACGATTGTCCGACGTAT | TCCGCCCAGGGACGAATACAGAGAA | ta | GAAGAGTCTTCCTTTACG |
| **14** | GAGGAGGGCAGCAAACGG | aa | GCGAAGTGACGATGATGACCTGTGT | TCGGGTCCGAATAACGTTGGATGAT | ta | GAAGAGTCTTCCTTTACG |
| **15** | GAGGAGGGCAGCAAACGG | aa | AAGTCGTGCAAGATCTTATCGACGT | TTTTCGACAGCGACGGATCGGTTTG | ta | GAAGAGTCTTCCTTTACG |
| **16** | GAGGAGGGCAGCAAACGG | aa | CAGATCCGACGATGACGGATTCTGT | CGCACCGCAATACCCTGCTGTAGTA | ta | GAAGAGTCTTCCTTTACG |
| **17** | GAGGAGGGCAGCAAACGG | aa | GATGATGGTGACGATGACATCGTTA | CAAGAGTGATTGTCGCAACACGACG | ta | GAAGAGTCTTCCTTTACG |
| **18** | GAGGAGGGCAGCAAACGG | aa | GTGAAGTTGGAGTATGATAACCGAA | TCATCTGGGTACCTGGAGATAGAAA | ta | GAAGAGTCTTCCTTTACG |
| **19** | GAGGAGGGCAGCAAACGG | aa | ACGATTTCGACTGAACGCAATTTAT | CGCGAAAGACATTATCTCGATGATC | ta | GAAGAGTCTTCCTTTACG |
| **20** | GAGGAGGGCAGCAAACGG | aa | CTTGTAATCGAAACCATTATGTCAG | CTCTAACGTAAAATACCCGCCGATG | ta | GAAGAGTCTTCCTTTACG |

**Table S8.** Probe pairs designed for *Pycnogonum litorale Sp6-9* HCR *in situ* hybridization (B2 initiator).

| Pair | Initiator | Spacer | Hybridization | Hybridization | Spacer | Initiator |
| --- | --- | --- | --- | --- | --- | --- |
| 1 | CCTCGTAAATCCTCATCA | aa | GCGTACTAACTGATGGCGGTTCACT | GTGTTCCCGGACTATGAATAGACGA | aa | ATCATCCAGTAAACCGCC |
| 2 | CCTCGTAAATCCTCATCA | aa | TTGCCGGCGATCATGCTGTGAGTTT | TCCGAATCACTACAACTGCTCGTCT | aa | ATCATCCAGTAAACCGCC |
| 3 | CCTCGTAAATCCTCATCA | aa | AAACGTTTGTTACAGACCGGACAAG | TGTTTGGCCAGATGGTCGGATCTCA | aa | ATCATCCAGTAAACCGCC |
| 4 | CCTCGTAAATCCTCATCA | aa | TCGTCGGAACGAGTGAAGCGTTTTC | GTGTGAGTACGCAGATGTCGCTGAA | aa | ATCATCCAGTAAACCGCC |
| 5 | CCTCGTAAATCCTCATCA | aa | CCGGTATGCCACCGTAAATGAGCTT | AACCAATTACACACGAACGGCCTCT | aa | ATCATCCAGTAAACCGCC |
| 6 | CCTCGTAAATCCTCATCA | aa | CATCCGGGAATGTGACAGCTGTGTA | TGAGACGTCTTGTTGTAAACCTTTC | aa | ATCATCCAGTAAACCGCC |
| 7 | CCTCGTAAATCCTCATCA | aa | GCCAGTGTACCTCCGTTGAGATCTT | ACAATTCGGACAATCGCAAGTCGCT | aa | ATCATCCAGTAAACCGCC |
| 8 | CCTCGTAAATCCTCATCA | aa | TATCTTGCAGAATATGTTGACCCGA | CAGCCGACGGCAACATCGTCTTATA | aa | ATCATCCAGTAAACCGCC |
| 9 | CCTCGTAAATCCTCATCA | aa | TGTATGTCCGAGTCCATAATCGCCA | TAATAAAGGTCCAGCTGCAGCTGCG | aa | ATCATCCAGTAAACCGCC |
| 10 | CCTCGTAAATCCTCATCA | aa | AAAACCAGGCATTGCCGTTGCCGAA | TGCAGTCGGATAACTTAACTGAGAA | aa | ATCATCCAGTAAACCGCC |
| 11 | CCTCGTAAATCCTCATCA | aa | CGACGTGCTATTCAACGTATTGACA | AGCATGAGAGTGCATGTCCCACCAT | aa | ATCATCCAGTAAACCGCC |
| 12 | CCTCGTAAATCCTCATCA | aa | GCCACGTTCATATTGAAAGGCCAAC | GTGACTTCTGGTTTAACCGTCGCTG | aa | ATCATCCAGTAAACCGCC |
| 13 | CCTCGTAAATCCTCATCA | aa | CTCGAGAATACATCGTTCCAGCTGA | CGTATGGATGATGATGAGCTGCTGC | aa | ATCATCCAGTAAACCGCC |
| 14 | CCTCGTAAATCCTCATCA | aa | AGCTTGAGCACTTAAGCTTCCAGCA | GTGGGCCGAACCGCTATGATTATTC | aa | ATCATCCAGTAAACCGCC |
| 15 | CCTCGTAAATCCTCATCA | aa | AACTGACAGGACTGGACGAGACAGA | AGATATCCGAATTACTACCATAGGC | aa | ATCATCCAGTAAACCGCC |
| 16 | CCTCGTAAATCCTCATCA | aa | AACCACTCTGATAATGCTGATTAGA | TGCCTCTTGCATTTGTCACCGCACA | aa | ATCATCCAGTAAACCGCC |
| 17 | CCTCGTAAATCCTCATCA | aa | GTTGTTCGTCGGATTCAGAGTGACA | GCTATGATGATGATGATGATGATTG | aa | ATCATCCAGTAAACCGCC |
| 18 | CCTCGTAAATCCTCATCA | aa | ACCCGTCGGCGACATCACATTGTTA | CGTAGAATCGGAAGTCGTCCTATTT | aa | ATCATCCAGTAAACCGCC |
| 19 | CCTCGTAAATCCTCATCA | aa | GTTGTCGGTGGACTAATCGACGGTT | CTCAGATGAGGAGACGTTGCAATGG | aa | ATCATCCAGTAAACCGCC |
| 20 | CCTCGTAAATCCTCATCA | aa | ATCGGCTAACGGAGGAGGACTCTTA | AGGATGGAATCCTTTGCCGACTGCT | aa | ATCATCCAGTAAACCGCC |
| 21 | CCTCGTAAATCCTCATCA | aa | ATTGCCAAAGGTGTATTCCCTAAAC | GCTAATTTATTACACTGAGCCGCCA | aa | ATCATCCAGTAAACCGCC |

**Table S9.** Primer pairs designed for *Phalangium opilio Lim1* RNAi experiments.

| **Primer Name** | **Primer Sequence** | **Amplicon Length** |
| --- | --- | --- |
| Popi_Lim1_for | GGCCGCGGCCTCATCACTCCCACCATTC | 665 bp |
| Popi_Lim1_rev | CCCGGGGCAAACTGTCGTTTTCCGGTTG |  |
